# Benchmarking of spatial transcriptomics platforms across six cancer types

**DOI:** 10.1101/2024.05.21.593407

**Authors:** Sergi Cervilla, Daniela Grases, Elena Perez, Francisco X. Real, Eva Musulen, Julieta Aprea, Manel Esteller, Eduard Porta-Pardo

## Abstract

Spatial biology experiments integrate the molecular and histological landscape of tissues to provide a previously inaccessible view of tissue biology, unlocking the architecture of complex multicellular tissues. Within spatial biology, spatial transcriptomics platforms are among the most advanced, allowing researchers to characterize the expression of thousands of genes across space. These new technologies are transforming our understanding of how cells are organized in space and communicate with each other to determine emergent phenotypes. This is particularly important in cancer research, as tumor evolution is shaped not only by the genetic properties of cancer cells but also by how they interact with the tumor microenvironment and their spatial organization. While many platforms can generate spatial transcriptomics profiles, it is still unclear in which context each platform better suits the needs of its users. Here we compare the results obtained using 5 different spatial transcriptomics (VISIUM, VISIUM CytAssist, VisiumHD, Xenium, and CosMx) and one spatial proteomics (VISIUM CytAssist) platforms across serial sections of 6 FFPE samples from primary human tumors covering some of the most common forms of the disease (lung, breast, colorectal, bladder, lymphoma and ovary). We observed that the VISIUM platform with CytAssist chemistry yielded superior data quality than manual VISIUM. On the other hand, Xenium consistently produced more reliable results for in situ platforms, with better gene clustering and fewer false positives than CosMx. Importantly, these platform-based variations didn’t significantly affect cell type identification. VisiumHD offers the best options of both worlds, with subcellular resolution and whole-transcriptome coverage, albeit with some limitations that need to be accounted for in downstream analyses. Finally, by comparing VISIUM protein profiles with the spatial transcriptomics data from all four platforms on each sample, we identified several genes with mismatched RNA and protein expression patterns, highlighting the importance of multi-omics profiling to reveal the true biology of human tumors.

## Introduction

Personalized cancer care relies on the molecular and cellular characterization of individual tumors. The field took a significant step further thanks to the analysis of bulk molecular profiles of tumors, including large-scale projects such as The Cancer Genome Atlas^1^, the International Cancer Genome Consortium (ICGC)^2^, or the Clinical Proteogenomics Tumor Analysis Consortium^3^. These endeavors have paved the way for a deeper understanding of critical oncogenic processes, including identifying cancer driver genes^4,5^, the interplay between molecular alterations in cancer cells and their surrounding tumor microenvironment^6,7^, or catalogs of clinically actionable mutations^8,9^. However, the technological limitations of bulk molecular characterization made it impossible, at the time, the in-depth analysis of the rich and complex ecosystem of the different cell types that co-exist within a tumor.

The first steps in that direction came from the analysis of tumors with single-cell technologies. Thanks to single-cell -omics, we are no longer limited to studying the average signal from a tumor sample; instead, we can now disaggregate it into its individual cells and study them separately. This revealed a breadth of mutational^10^, epigenetic^11^, and transcriptional^12^ heterogeneity in cancer cells and the TME.

While a significant step forward, single-cell technologies still have a significant limitation: due to the disaggregation of the tumor, all the information regarding the spatial location of the cell in its original tissue is lost. In this context, spatial “-omics” platforms provide a significant advantage over single-cell approaches^13^. Powered by technological advances that quantify the spatial distribution of thousands of molecules, we can now study the spatial distribution of DNA alterations^14^, epigenetic profiles^15^, or the expression of genes^16–23^ and proteins^24–26^.

Within spatial omics, spatial transcriptomics is currently probably the most widely used spatial technology in cancer research. This is, among others, thanks to the availability of multiple commercial platforms. These can be classified into two groups depending on the readout used to quantify gene expression. On the one hand, we have sequencing-based technologies such as VISIUM^16^, or SlideSeq^22^. These rely on the use of nucleic acid tags to identify the location of the molecule within the slide. On the other hand, other platforms (such as CosMx^20^, Xenium^23^ or MERSCOPE^18^) rely on *in situ* imaging of the RNA molecules, which are identified with fluorescent probes.

Given the variety of chemistries and platforms available for spatial transcriptomics, it is important to understand the advantages and limitations of the different options so researchers can design their experiments accordingly. Here, we compare the results of five spatial transcriptomics (VISIUM manual, VISIUM CytAssist, VISIUM-HD, Xenium, and CosMx) and one spatial proteomics (VISIUM CytAssist) platforms when profiling serial FFPE sections from six human primary untreated tumors: breast, lung, colorectal, ovary, bladder, and lymphoma.

Our results show several consistent patterns across all studied cancer types. For example, the inclusion of CytAssist and the new chemistry has significantly improved the quality of the data of the VISIUM platform. Regarding the *in situ* platforms, Xenium seems to consistently yield more reliable data than CosMx, with a large fraction of genes having higher spatial clustering in the former than in the latter, while also having a lower fraction of false positives. Remarkably, though, this is not translated into major differences in cell type annotations between both platforms. A newer version of VISIUM, VISIUM-HD, offers whole-transcriptome coverage at subcellular resolution. Finally, by correlating the VISIUM-protein profile with the spatial transcriptomics of all four platforms of each sample, we were able to identify several genes that diverge in the spatial expression of their RNA and associated proteins, highlighting the importance of multi-omics profiling of biomedical samples. Overall, our results highlight several important differences across spatial -omics platforms that users need to be aware of when designing their experiments.

## Results

### Generation of matching spatial transcriptomics and proteomics data

We generated spatial transcriptomics data (VISIUM manual, VISIUM CytAssist, VisiumHD, Xenium, and CosMx) from serial sections of six primary untreated tumor samples covering some the most common cancer types: colorectal (adenocarcinoma), breast (infiltrating carcinoma of non-special type), lung (acinar invasive adenocarcinoma), lymphoma (diffuse large B-cell lymphoma), ovarian (high-grade serous carcinoma), and bladder (muscle-invasive bladder cancer) (**Figure 1a**). We analyzed consecutive sections of formalin-fixed, paraffin-embedded (FFPE) tumors obtained on the same day for all platforms, except those for VISIUM manual and VisiumHD, which were obtained at different time points and between 25 and 50 μm difference in the z-axis from the block (**Figure 1b,c**).

**Figure 1.**
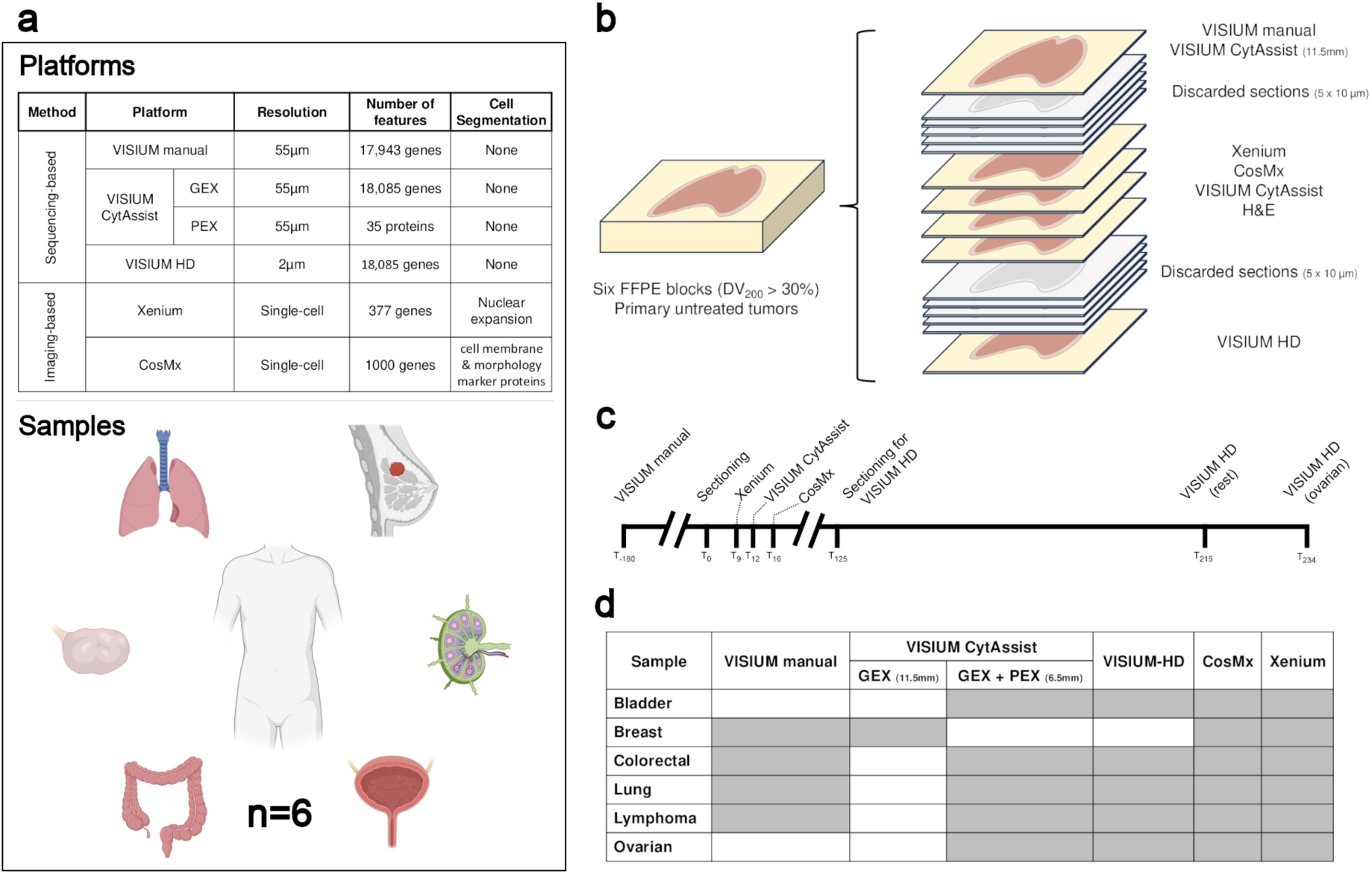
Experimental design. **a)** Summary of platforms and tissue types used in the benchmark. **b)** Schematic representation of the tissue section collected in a FFPE block for each of the six tumor types. **c)** Timeline showing the generation of the different datasets. **d)** Table of the samples generated for each technology used in the benchmarking.

All experiments successfully yielded spatial transcriptomics profiles (**Figure 1d**), except for the VISIUM CytAssist breast sample which partially detached during the 95°C decrosslinking step of the protocol. This led to the tissue folding on top of itself and, therefore, an unusable library. However, we had generated VISIUM CytAssist data from the same sample using the 11.5 mm capture area, which we used as a replacement for the analyses presented here. Additionally, we generated spatial proteomics data using the VISIUM CytAssist immuno-oncology antibody panel (n = 35 proteins). In this case, this data is not available for the breast cancer sample for the reasons explained above. Overall, after filtering low-quality spots and cells (**Methods**), we captured 18,672 spots with VISIUM manual, 33,117 spots with VISIUM CytAssist, 2,551,540 8×8 µm bins with VisiumHD, 667,916 cells with CosMx, and 2,645,127 cells with Xenium.

### Whole-transcriptome coverage at a multicellular resolution

Sequencing-based spatial transcriptomics platforms have become a popular choice for researchers, as they provide whole-transcriptome coverage at near single-cell resolution. Among these methods, VISIUM from 10x Genomics stands as a leading technology, offering robust spatial transcriptomics through spatially barcoded capture arrays. There are, so far, three iterations of the technology: VISIUM manual, VISIUM CytAssist, and VISIUM-HD. We first focus on the comparison of the first two, as they both offer 55µm resolution, and represent the first iterations of the technology as well as most of the data generated with this platform so far. The main differences between both include how one places the tissue section on the VISIUM slide (manually or with the help of a CytAssist tool), as well as differences in the number of pairs of probes per coding gene (one in the manual compared to three in the CytAssist). According to 10x Genomics, the higher number of probes per gene, allows the analysis of samples that have more degraded RNA. While 10x recommends a DV200 over 50% for the manual version, the recommended threshold for the CytAssist version is >30%. Also, according to 10x Genomics, the use of the CytAssist to run the protocol leads to lower lateral diffusion of the RNA molecules during the protocol. To test the strengths and weaknesses of each VISIUM version, we generated data from the same samples with both platforms.

Our results show that, overall, the vast majority of genes (∼15,000) are consistently detected using both chemistries (**Figure 2a**), however, CytAssist tends to have more uniquely detected genes (∼2,000) than VISIUM manual (∼200), consistent with the fact that VISIUM CytAssist has three probes per gene instead of only one. Similarly, the number of detected genes (**Figure 2b**) and UMI (**Figure S2**) per spot also tends to be higher in VISIUM CytAssist than in the VISIUM manual.

**Figure 2.**
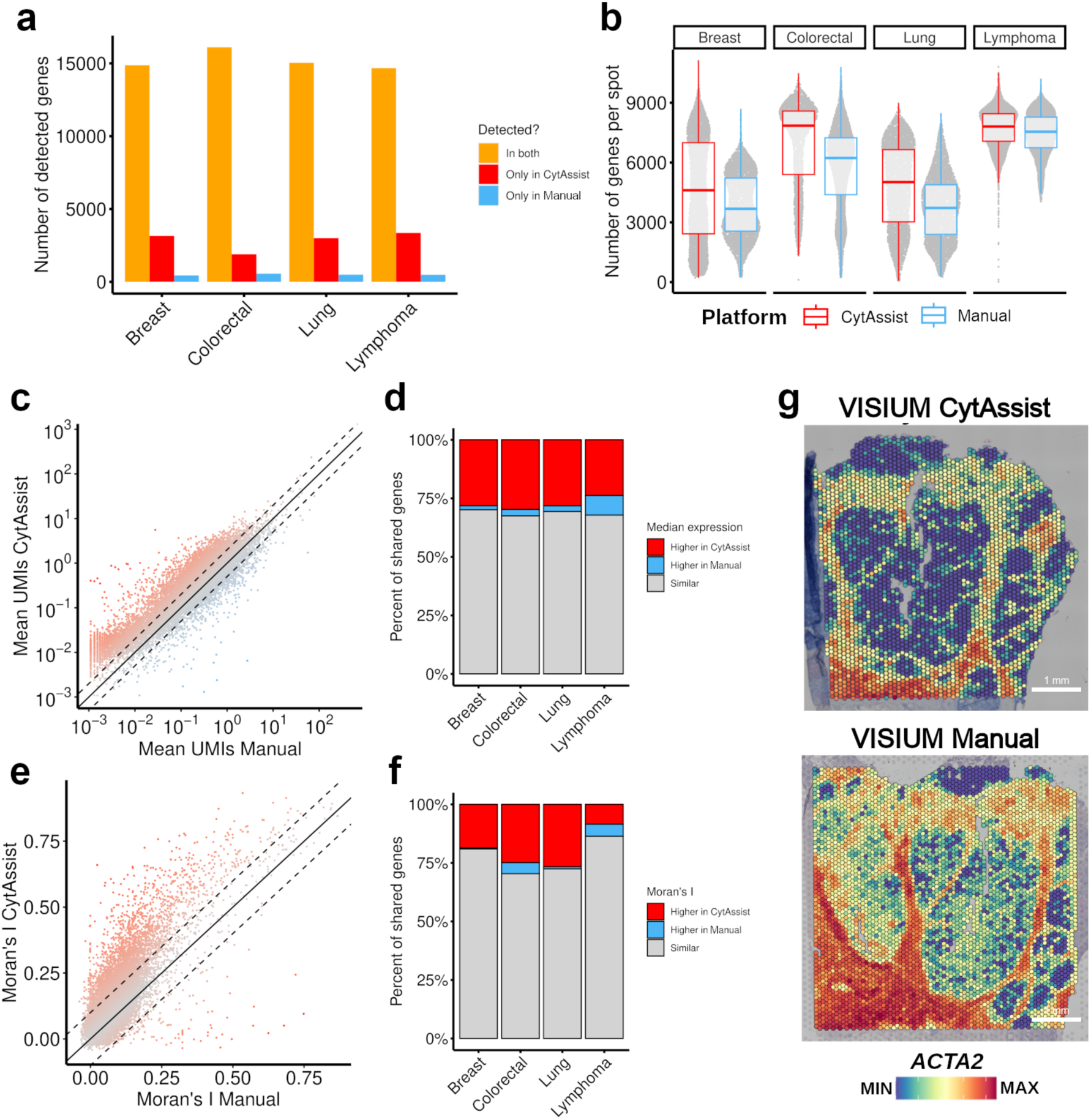
Comparison of VISIUM manual and CytAssist. **a)** Barplot of the number of detected genes in each sequenced-based platform colored by shared and unique genes. **b)** Distribution of the number of unique genes detected in each spot across samples and platforms. **c)** Scatter plot of mean UMI per genes in VISIUM manual and CytAssist in the lung sample. Color intensity of dots represents the distance from the diagonal (equal UMI detection across platforms). **d)** Barplot showing the relative number of genes that have similar numbers of transcripts or are higher in one platform for each sample. **e)** Scatter plot of global Moran’s I for each gene in VISIUM and CytAssist in the lung sample. Color intensity of dots represents the distance from the diagonal (equal UMI detection across platforms). **f)** Barplot showing the relative number of genes that have similar spatial distribution or are higher in one platform for each sample. **g)** Gene expression of *ACTA2* of the colorectal sample using CytAssist (top) and VISIUM (bottom).

To assess how these differences affect the results of individual genes, we calculated the average UMIs per gene in all spots of each sample in each platform. As we show for the lung sample (**Figure 2c, Figure S3**), while the correlation between VISIUM manual and CytAssist is very high, there is a significant fraction of genes with higher UMI counts when using the CytAssist platform, particularly at the lower range of expression (mean UMI per spot below 0.01). Overall, across all samples, between 25% and 30% of genes had twice as many mean UMIs or more in VISIUM CytAssist than in VISIUM manual (**Figure 2d**), whereas only between 2% and 6% had the opposite pattern.

Finally, to quantify to what extent these differences in gene detection could impact on the overall data quality, we used Moran’s I to quantify the randomness of the spatial distribution of the expression of each gene. The rationale is that, since tissues are spatially structured and many genes tend to be cell type specific, most genes should have a non-random spatial distribution (i.e. higher Moran’s I values). Using again the lung sample as an example (**Figure 2e, Figure S4**), one can see that, while there is an overall good correlation between the Moran’s I of each gene obtained by both platforms, many genes have higher Moran’s I with the CytAssist data. Putting the threshold at a Moran’s I 0.1 value higher in one of the two platforms, only between 0.5% to 7% of genes have a higher value in VISIUM manual, whereas between 19% to 27% do for the VISIUM CytAssist (**Figure 2f**).

This higher clustering with the CytAssist version of VISIUM can be seen in the expression pattern of *ACTA2* in the colorectal cancer sample (**Figure 2g**). This gene, which is associated with the presence of fibroblasts, overlaps the fibrotic area of the sample in both platforms, but the non-fibrotic area in the case of manual VISIUM also shows some degree of expression, suggesting a noisier pattern with potential false positives. Overall, our results show that the use of VISIUM CytAssist, along with the changes in the chemistry of the reagents, have led to a significant improvement in the data generated by VISIUM manual.

### Subcellular resolution with limited gene panel size: *in situ*

An emerging alternative to sequencing-based spatial transcriptomics platforms are those based on *in situ* transcript quantification through imaging. These rely on fluorescent probes and high-resolution imaging to capture the precise location of individual mRNA molecules at subcellular resolution, which are then assigned to individual cells using cell segmentation approaches. There are currently a variety of platforms available, but here we focused on two of the more widely-used: CosMx (from Nanostring) and Xenium (from 10x Genomics).

While the general concept of both platforms is similar, there are important differences that need to be considered before any comparison can be made. For example, neither platform generates (yet) whole-transcriptome data but, instead, they focus on gene panels. In the case of Xenium, we used the human multi-tissue panel which includes 377 genes, whereas for CosMx we used the 1000 genes panel. Both platforms also differ in the details of how they acquire the data. While Xenium can scan a continuous area of 1.1 cm by 2.4 cm, CosMx scans several user-defined fields-of-view (or FOV) that are 0.5mm by 0.5mm. In our case, we scanned an average of 58 FOVs per sample with CosMx (totaling 87.5 mm^2^ for all samples), whereas we scanned 394.7 mm^2^ with the Xenium. Finally, another important difference is how both platforms segment the cells that they capture. In the case of Xenium, it uses DAPI to identify the nucleus and then expands the perimeter to identify the solution that best limits the space occupied by each cell. On the other hand, CosMx relies on both, DAPI staining for the nucleus, as well as membrane staining, identifying both cell compartments.

These technical differences were reflected in the data itself. For example, given the larger area scanned by Xenium, we captured significantly more cells for each sample than with the CosMx (**Figure 3a**). On the same line, CosMx has an important artifact on its FOVs, where cells that are at its border have an order of magnitude less counts than cells in the middle of the FOV (**Figure 3b**). Finally, regarding the impact of the gene panel size, since the CosMx gene panel has significantly more coverage than the Xenium multi-tissue that we used (1000 genes compared to 377), we detected between 2 and 3 times more genes and transcripts (counts) per cell with the CosMx than with the Xenium (**Figure 3c,d**). To limit the impact of all these differences in the downstream comparisons, unless stated otherwise, the rest of the experiments exclude CosMx cells at the border of the FOV, are limited to cells in areas shared between both platforms (**Figure 3e, Figure S5**), and only include genes shared between both panels (n = 125).

**Figure 3.**
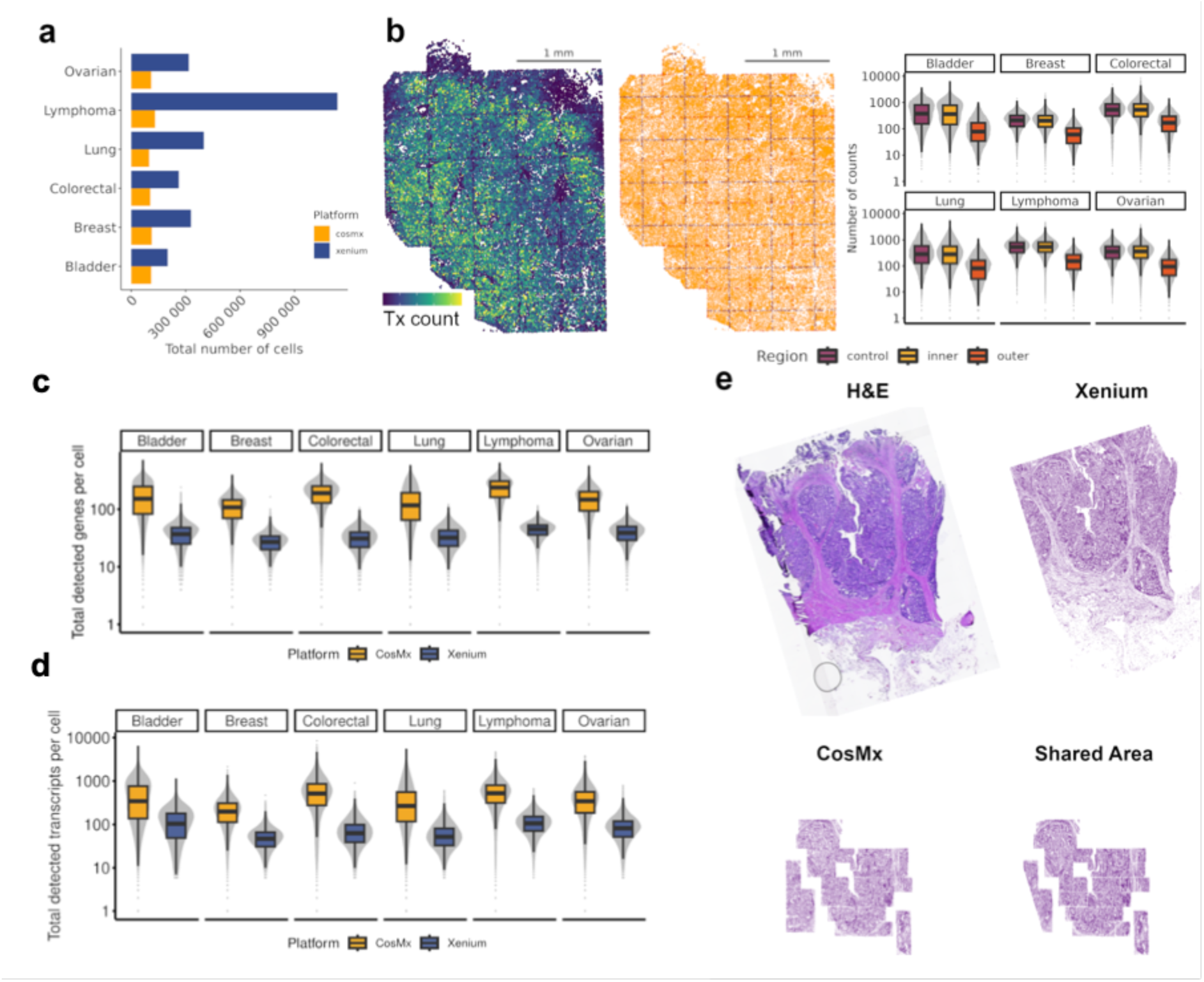
Main differences between *in situ* platforms. **a)** Barplot of the number of detected cells in each image-based platform (whole scanned area). **b)** The left plot shows the number of transcripts detected per cell in a small area of the bladder sample in CosMx. The middle plot shows the cells in the bladder sample (CosMx) colored according to their distance to the FOV edge: in red the control (between 2000 and 2100 px), in orange the outer area (36px from the edge), and the remaining cells in yellow. The right boxplot shows the number transcripts detected in each cell and in each CosMx sample. **c)** Boxplot comparing the number of unique genes detected per cell in CosMx and Xenium across six tumor samples. **d)** Boxplot comparing the number of transcripts detected per cell in CosMx and Xenium across six tumor samples. **e)** Representation of the colorectal tissue analyzed and compared: hematoxylin and eosin from a section of the FFPE block (upper, left), cells captured using Xenium (upper, right), cells captured using CosMx (bottom, left) and cells that are in the shared regions between Xenium and CosMx.

The number of cells in the shared areas of each sample converged toward similar values, even if Xenium still has an average of 5% to 30% more cells (**Figure 4a**). Given the differences in staining and segmentation, we wondered whether this difference in cell numbers could be explained by these factors. In that sense, we observed that cells analyzed with Xenium were, on average, bigger than in CosMx in all samples but lymphoma (**Figure 4b**).

**Figure 4.**
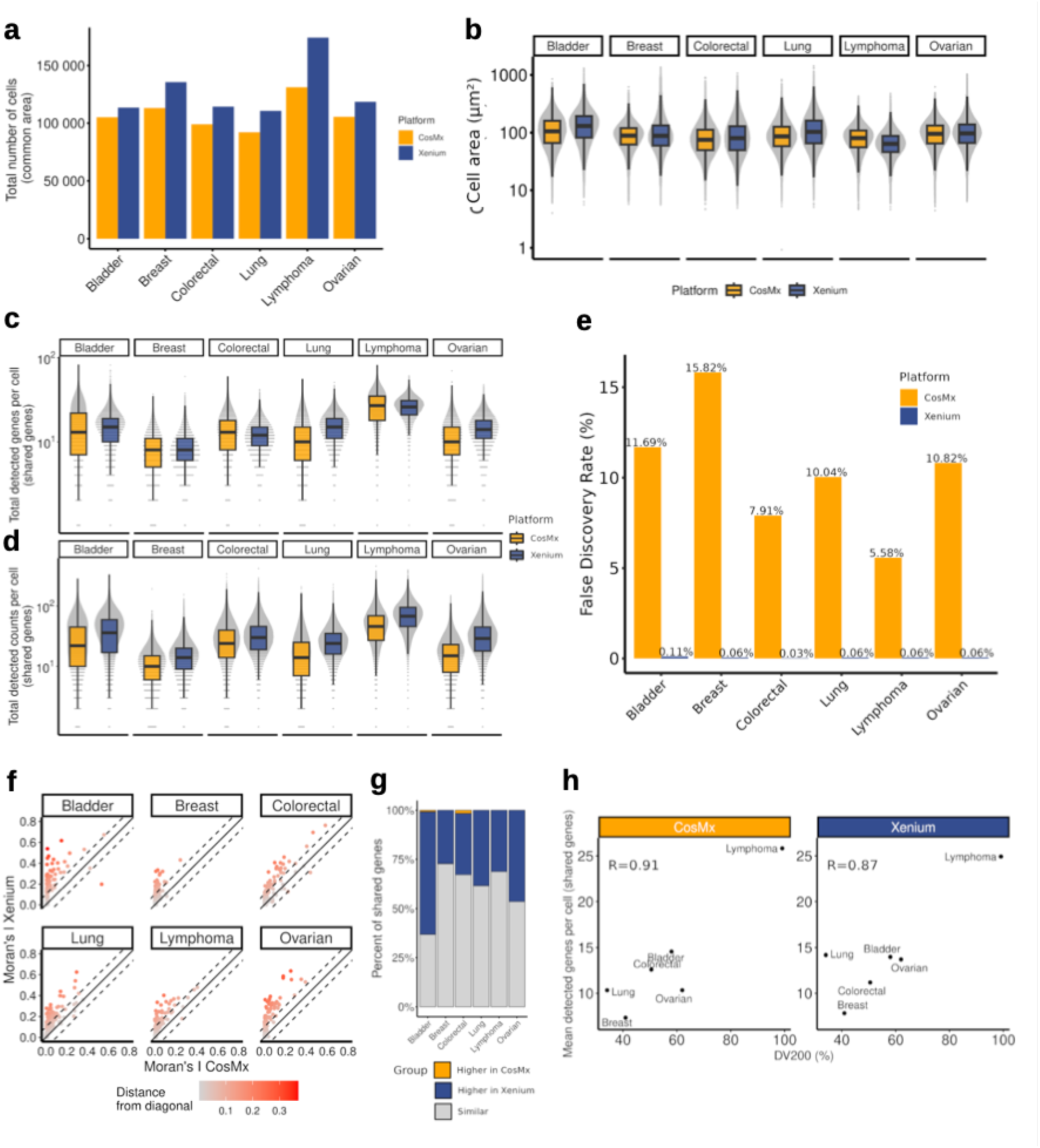
Comparison of *in situ* platforms after accounting for the main differences. **a)** Barplot of the number of cells within the shared regions between CosMx and Xenium in the different tumor types. **b)** Boxplot comparing the distribution of the cell area (µm²) in CosMx and Xenium. **c)** Boxplot of the number of unique genes detected per cell (y-axis, in log scale) in CosMx and Xenium considering only the 125 genes shared between the two gene panels. **d)** Boxplot of the number of transcripts detected per cell (y-axis, in log scale) in CosMx and Xenium considering only the 125 genes shared between the two gene panels. **e)** Barplot of the False Discovery Rate in CosMx and Xenium in each sample. **f)** Scatter plot comparing the spatial distribution (Moran’s I) of shared genes in CosMx and Xenium. The dots are colored according to the distance to the diagonal (equal Moran’s I across platforms). **g)** Barplot showing the relative number of genes with similar spatial distribution or higher in one of the platforms in each sample. **h)** Correlation between the DV200 of each sample (x-axis) and the mean detected genes per cell (y-axis) in both platforms.

We next looked at the values for shared genes and transcripts across both platforms. The average number of genes detected per cell was very similar (around 13 in CosMx and 14 in Xenium), with Xenium being slightly higher in bladder, lung, and ovarian and CosMx in colorectal (**Figure 4c**). In terms of gene counts per cell, however, Xenium gave a higher output (around 26 in CosMx and 37 in Xenium) in all samples (**Figure 4d**).

Next, we calculated the false discovery rate (FDR) for both platforms. Given the lack of ground truth, calculating the FDR can be challenging. Furthermore, CosMx and Xenium have different numbers of control probes (10 in CosMx and 20 in Xenium). For this reason, we calculated the FDR value assuming that all counts are true positives, but focusing only on the 125 shared genes. This showed that CosMx has an FDR around 2%, whereas Xenium only 0.01%, even if the latter has twice as many negative probes (**Figure 4e**). In the same line, the difference in counts between negative probes and gene probes was significantly higher in Xenium than in CosMx (**Figure S6**). In CosMx, gene probes had around 3 times as many counts as negative probes, whereas in Xenium the ratio was around 450.

Following the same reasoning as with the sequencing-based platforms, we calculated the Moran’s I of each gene in each platform to assess the spatial clustering of the signal detected by each platform. These results showed that Moran’s I of Xenium data was higher than for CosMx in ∼40% of the genes (**Figure 4f,g**). Finally, we explored the relationship between the quality of the RNA in the sample, measured as DV200, and the mean detected genes per cell. Overall, we observe a strong correlation between both variables in both platforms, suggesting that users should assess the RNA quality of their samples before proceeding to the experiment (**Figure 4h**).

Given the differences in panel size, sensitivity, false discovery rate, and cell segmentation, we wondered to what extent both *in situ* platforms would yield similar insights into the biology of the samples. We first annotated all cells from both platforms with their estimated cell type based on their expression profile using SingleR^27^ (**Figure S7, Figure S8**). Next, we re-assessed the same metrics as in the previous section but for each cell type. Overall, we observed similar patterns as when analyzing the entire sample together. However, we found that cell area, number of unique genes and transcripts captured had specific patterns regarding the cell type. For example, cells residing in lower density regions such as smooth muscle and fibroblast had consistently higher cell size in Xenium, reflecting the impact of using nuclear expansion as cell segmentation strategy (**Figure 5a, Figure S9**). The only exceptions were T and NK cells in the lymphoma sample, which could be related to the higher differences of cells found and the lower overall cell size for Xenium. Regarding gene and transcript sensitivity, substantial differences were noted between platforms, particularly within immune cells. Notably, only cancer cells of epithelial origin demonstrated robust characterization, with no significant differences observed across these metrics.

**Figure 5.**
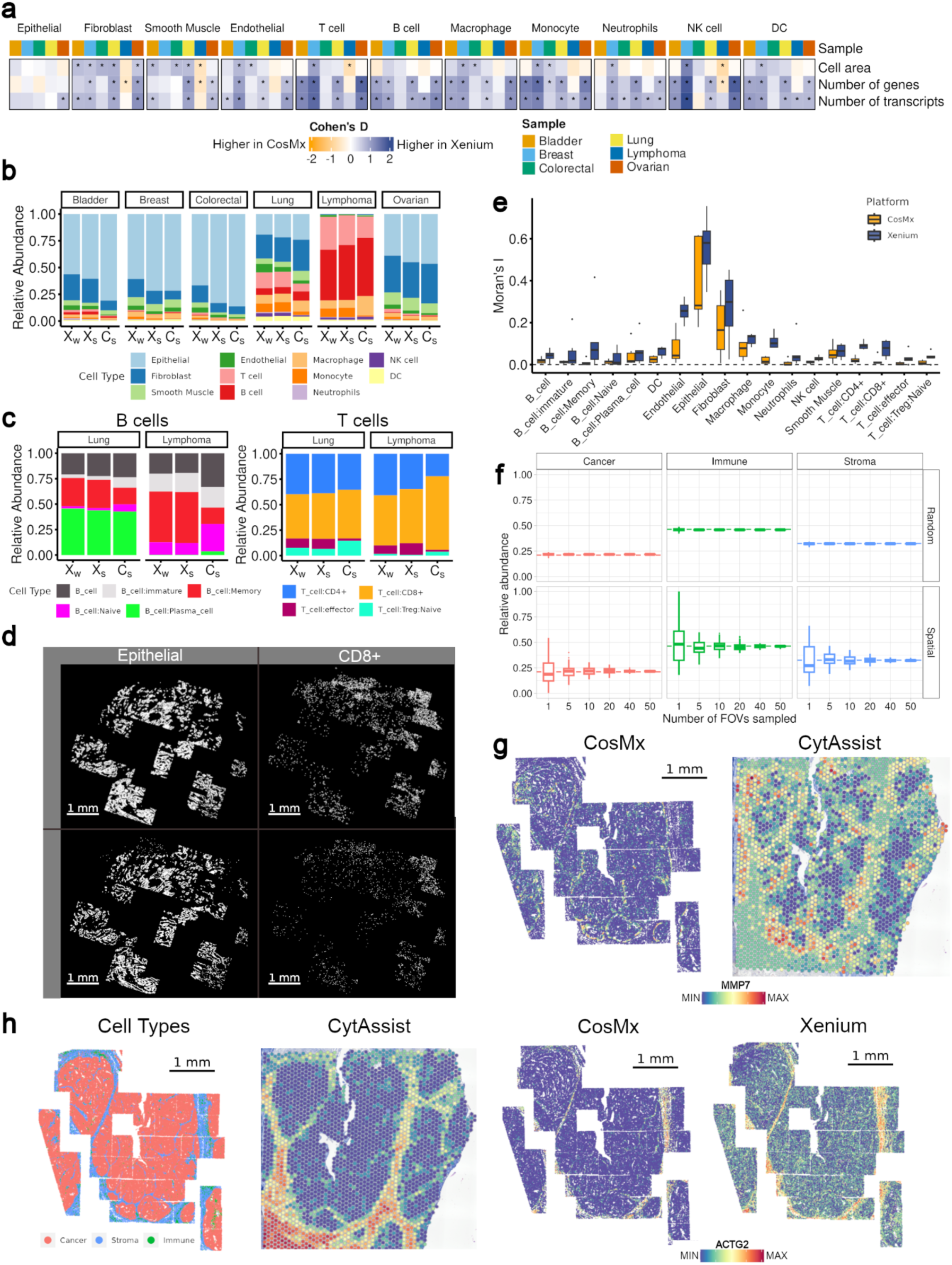
Identifying different cell types with each *in situ* platform. **a)** Differences by cell type between Xenium and CosMx in cell size (top row), number of genes per cell (middle row), and number of transcripts per cell (bottom row), across all samples. Only the 125 genes shared between the two gene panels were considered. **b)** Barplot showing the cell type relative abundance found in the whole Xenium sample (left), the common area of the Xenium cells (middle) and the common area of the CosMx cells in each tumor sample. **c)** Similar to b) but showing the sub-type relative abundance within B cells (left) and T cells (right) for lung and lymphoma samples. **d)** Spatial distribution of epithelial cells (left) and CD8+ T cells (right) in a zoom area of the lung sample for Xenium (top) and CosMx (bottom). **e)** Boxplot showing the global Moran’s I computed in the binary data for each cell identity, sample and platform. **f)** Boxplot showing the effect of sampling smaller regions to compute relative abundance in the xenium lung sample, of Cancer (epithelial), Stroma cells (fibroblast, smooth muscle and endothelial) and Immune (remaining cell types). In the Random category, the subset was determined by randomly sampling across the Xenium (shared area), while in the Spatial category, the sampling was based on FOV as a sampling unit. In each group number of sampled FOV, we used 100 stochastic iterations to compute the abundance of each cell group. Dashed lines represent the relative abundance of the Xenium (shared area) sample. **g)** *MMP7* gene expression in the CosMx (left) and CytAssist (right) samples. **h)** Representation of the cancer, stromal and immune cells in the Xenium colon sample (first) and *ACTG2* gene expression in the CytAssist (second), CosMx (third) and Xenium (fourth) samples.

We also assessed the differences in cell type composition of each sample inferred from each platform. Despite the differences in gene panel size, counts per gene, and cell size, the estimated cell composition in the shared area of both platforms is very similar in all samples, except the bladder sample, in which CosMx assigns more epithelial cells and fewer stromal and immune cells than Xenium (**Figure 5b**). Additionally, we noted notable differences in the assignment of monocytes and macrophages. While the total sum of these cell types remained consistent across platforms, CosMx allocated a smaller fraction to monocytes. Another aspect that caught our attention is that the estimated cell fractions in a given sample using data from the same instrument can differ significantly depending on how much area of the sample one is studying. This can be seen in the Xenium data, for which we had two values: the cell fraction from the entire scanned area and that corresponding to the subset of the area shared with CosMx (**Figure 5b**). While these two values were relatively similar in four samples, there were significant differences in the other two (breast and colorectal). Lastly, we explored their performance in identifying immune subtypes. In the lung and lymphoma samples, which contained a higher proportion of immune cells, we found that the distributions of B and T cell subtypes were nearly identical, except for memory and naive B cells, as well as effector and regulatory T cells **(Figure 5c, Figure S10**).

Despite their similar captured cell type composition, we investigated whether these cells exhibited comparable spatial distributions, given our previous finding that CosMx transcripts are noisier. For instance, if we focus on the lung epithelial cells across space, we find a similar spatial pattern that resembles the tissue morphology in both platforms (**Figure 5d, left**). However, for CD8+ T cells, CosMx exhibits nosier patterns with a lack of clear spatial clustering compared to Xenium, even though both platforms showed enrichment of CD8+ T cells in the right area of the tissue (**Figure 5d, right**). We used Moran’s I metric to quantify this observation, treating cell identity as binary data (**Figure 5e**). Xenium demonstrated higher spatial clustering of cell types, with notable differences in endothelial cells, CD4+ T cells, CD8+ T cells, monocytes, and memory B cells.

The generation of biases in the estimated cell composition of a sample when studying only a small fraction of the tissue, is relatively intuitive and has been previously described in detail for other data modalities^28^. This is very important, as there is a tendency to limit spatial biology studies to relatively small sections of samples to optimize resources and the use of precious samples. Therefore, we decided to understand further and quantify the impact of the size of the scanned area when using *in situ* platforms. To that end, we simulated the estimated cell fraction if we had sampled with the Xenium different numbers of FOVs equivalent to the size of the CosMx instrument (0.5mm x 0.5mm). Then, we compared them to the actual value obtained with the full area, along with an additional control consisting of a random sampling of the same number of individual cells that can be found on average in an FOV but from any region of the tumor, simulating a single-cell experiment. As expected, there is a correlation between the number of FOVs analyzed and how close the estimated cell composition resembles that of the entire tissue. For example, experiments sampling only one FOV, estimate the fraction of cancer cells anywhere between 5% and 55%, while the ground truth is 22% (**Figure 5f, Figure S11**). Notably, the variability in the simulated single-cell experiment was minimal and close to the ground truth, showing the relationship between spatial constraints and the number of cells needed to quantify cell abundance. Researchers must be aware of potential biases they might be introducing in their studies if they limit the amount of tissue being analyzed.

Another important aspect to consider is that the limited size of the gene panel in Xenium can lead to missing important biology that goes beyond cell type classification. For instance, the *MMP7* gene, the expression of which is associated with invasive tumor growth, distant metastasis, and chemotherapy resistance in colorectal cancer, is absent in the Xenium gene panel. In our colorectal sample, overexpression of *MMP7* was observed in the cancer cells at the immediate edge with the stromal cells (**Figure 5g**), facilitating the identification of epithelial cell subpopulations with potential clinical relevance. These biological insights would be lost due to the smaller size of the Xenium panel.

Overall, in the six samples we studied and with the panels and versions of the technology we have used, the raw data from Xenium seems to be, on average, of better quality than that of CosMx. These observations are consistent across most cell types present in our samples. That being said, CosMx data seemed to better capture the underlying biology for a subset of the genes. For example, the *ACTG2* expression, which encodes the gamma (γ)-2 actin protein, should be primarily found in smooth muscle cells. However, we observed gene expression within epithelial cells in Xenium across all samples, contrasting with the specificity to non-epithelial cells seen in CosMx, which we validated in the CytAssist data (**Figure 5h, Figure S12**).

### Comparison of VISIUM CytAssist and *in situ* platforms

The direct comparison of all four spatial transcriptomics platforms presented so far has some challenges. On one hand, both VISIUM versions are whole-transcriptome, whereas CosMx and Xenium are limited to 1000 and 377 genes, respectively. On the other hand, while the *in situ* platforms have single-cell resolution, both VISIUMs have a relatively low resolution of 55 μm, grouping between 5 to 15 cells (**Figure 6a**). For these reasons, certain metrics we have previously used such as Moran’s I, cannot be directly applied to this comparison. Furthermore, given the superior data quality from CytAssist when compared to manual VISIUM, in this section we focused only on the former.

**Figure 6.**
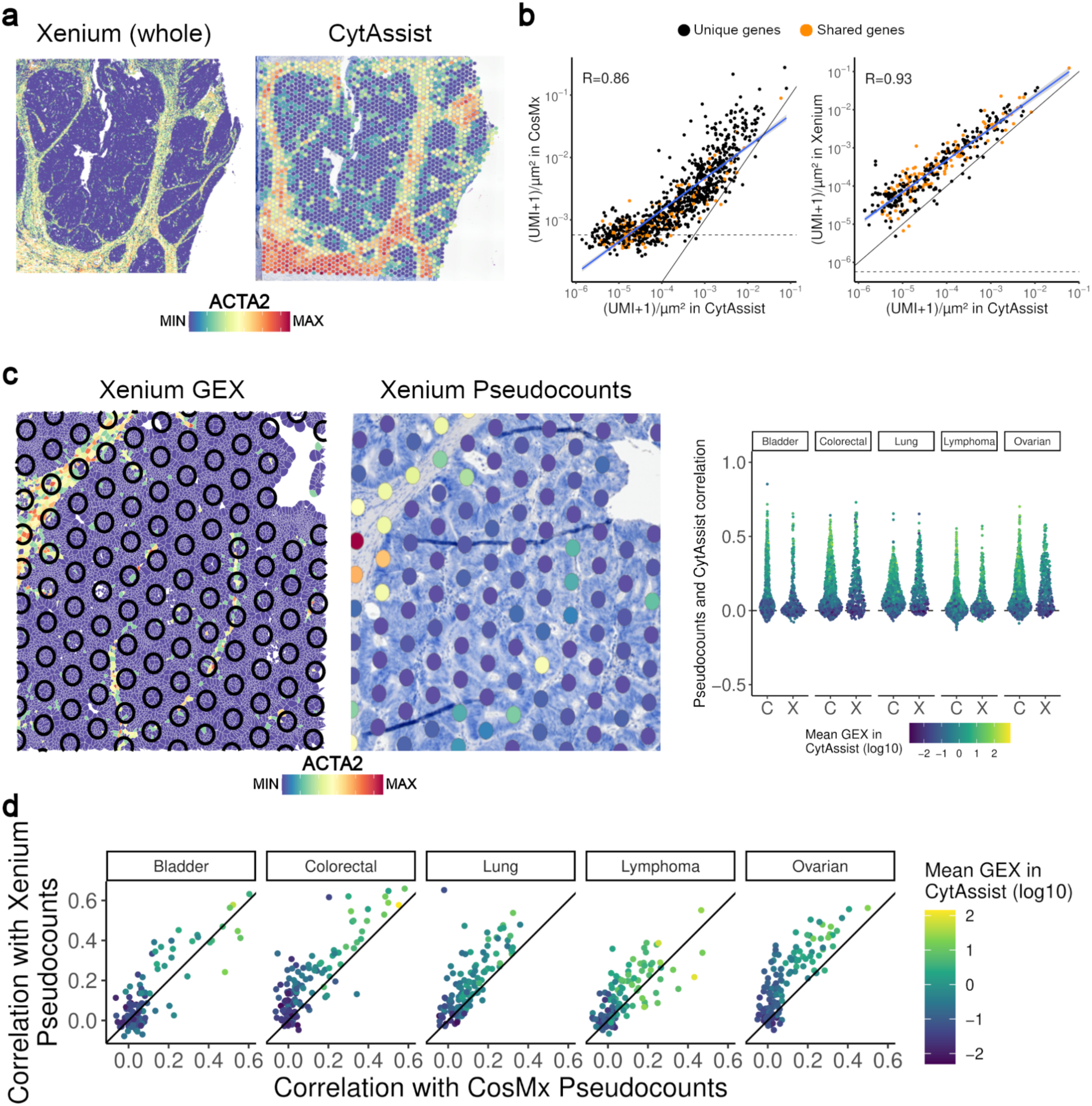
Comparison of VISIUM CytAssist, Xenium and CosMx. **a)** Plot showing the expression of *ACTA2* in serial sections of the colorectal sample with Xenium (left) and CytAssist (right). **b)** Scatterplots showing the correlation between density gene expression in CytAssist (x-axis) and CosMx (y-axis, left) or Xenium (y-axis, right). The horizontal dashed lines show the average detection rate of the negative probes in Xenium and CosMx. The diagonal dashed line shows the expected perfect correlation (i.e x = y). The blue line shows the correlation between CosMx/Xenium and CytAssist. **c)** Image showing the overlaid VISIUM spots in the Xenium sample (left) and the inferred pseudocounts if Xenium were used with the same resolution as Xenium for the same exact area (right). The right boxplot shows the distribution of correlation across space between actual CytAssist values and the CosMx (C) or Xenium (X) inferred pseudocounts (y-axis). Each dot represents a gene, and is colored depending on its mean gene expression in CytAssist. **d)** Scatterplot showing the relationship between the correlation of CytAssist and Xenium pseudocounts (y-axis) and CytAssist and CosMx pseudocounts (x-axis). Each dot is colored according to the mean expression of the gene in CytAssist.

We first compared the UMI density of each gene in the CytAssist data with each *in situ* platform (**Figure 6b, Figure S13**). The correlation between CytAssist and Xenium across all samples was very high (R = 0.93), whereas with CosMx was significantly lower (R = 0.86). This was largely driven by lowly expressed genes (UMI density per gene < 10^−3^), as they have an UMI density count virtually indistinguishable from the negative probes (horizontal dashed line in **Figure 6b**). This effect was consistent across all samples, and suggests that, while calls in lowly expressed genes in CosMx might not be as reliable, highly expressed genes probably do not have such problems.

To further dissect the differences between sequencing-based and *in situ*-based platforms, we aimed to quantify the spatial correlation between both. To do that, we aligned the location of the VISIUM CytAssist spots with the single-cell resolution image of both Xenium and CosMx. Next, we calculated the pseudo-bulk counts on each of the overlayed spots, giving us an approximate image of the *in situ* data if it had the VISIUM resolution (**Figure 6c**). Overall, the spatial expression correlation was, again, higher between Xenium and CytAssist (**Figure 6d**), particularly in lowly expressed genes. This was more pronounced in the colorectal and ovarian samples, and less so in the lung and lymphoma samples.

In conclusion, these results suggest that data from both types of platforms are relatively comparable, making it possible to leverage the strengths of each (single-cell resolution for *in situ* and whole-transcriptome characterization for sequencing-based) if one generates data from both from serial sections.

### Subcellular resolution and whole-transcriptome coverage

The four platforms presented so far require that the user makes a compromise between whole-transcriptome coverage at oligo-cell resolution (VISIUM, VISIUM CytAssist) and single-cell resolution limited to a panel of a few hundred genes (Xenium, CosMx). However, there is another platform that could potentially achieve both, single-cell resolution with whole-transcriptome coverage: VisiumHD.

Similar to the other VISIUM platforms, VisiumHD is sequencing-based, but instead of 55 µm circles with empty spaces between them, it is based on a continuous array of 2×2 µm square bins, each covered in uniquely barcoded oligonucleotides. Each capture area is a square of 6.5mm per side, yielding a total of ∼11M bins at the 2 µm resolution, which significantly increases computational demands. For this reason, other bin aggregations of larger sizes, such as 8 µm or 16 µm per side, are recommended, as they reduce the bin count to ∼700K and ∼160K, respectively. However, the choice of the bin size has an effect on the number of cells per bin as well as the number of bins per cell, which could have an impact on downstream analyses. To explore this phenomenon, we quantified two metrics: the frequency of multiple cells within a bin, and the average number of bins per cell. To do so, we overlapped a simulated binning grid of different sizes on the Xenium data of each sample (**Figure 7a**).

**Figure 7:**
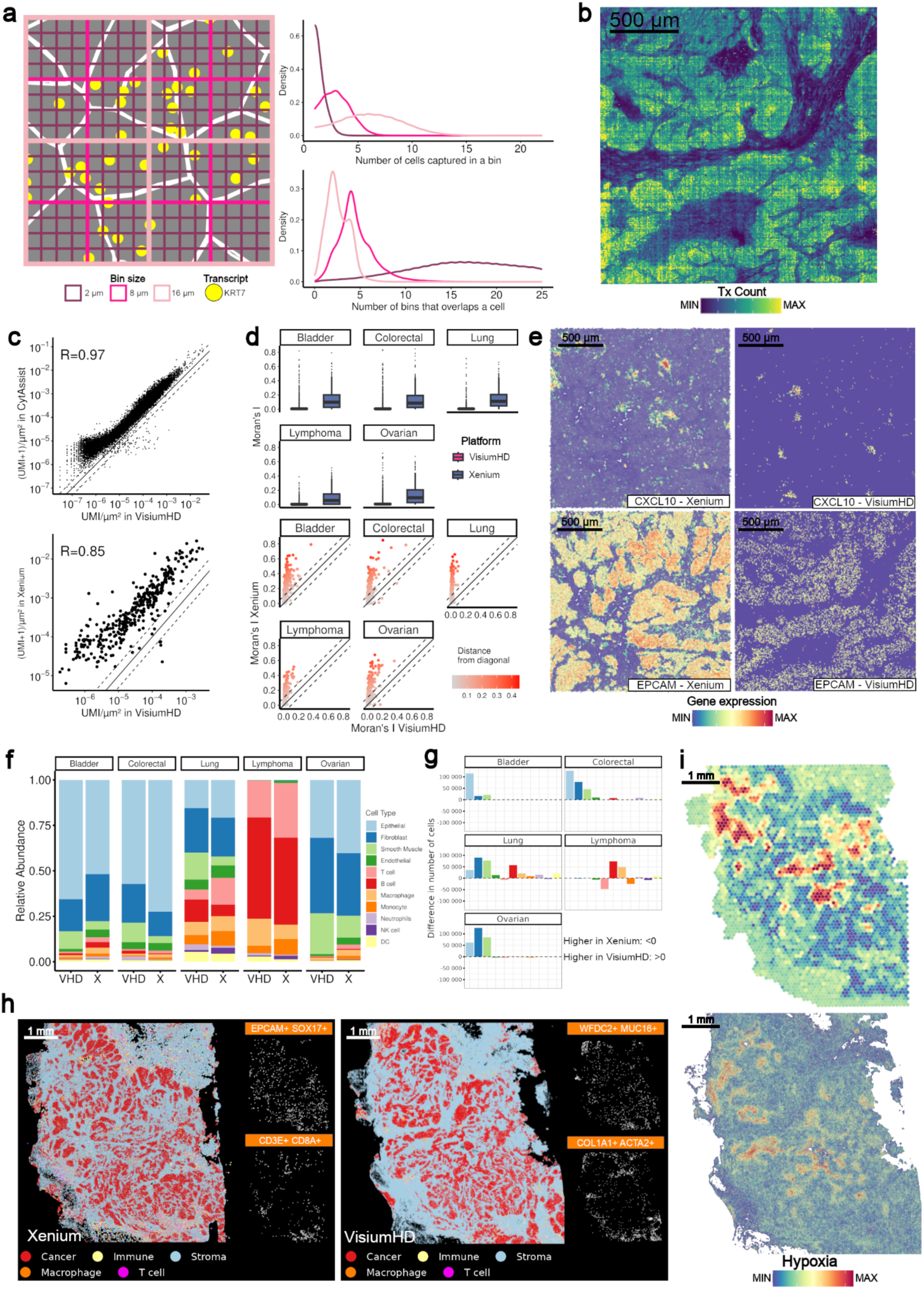
Comparing VisiumHD against VISIUM CytAssist and Xenium. **a) Left:** Small region (32×32 µm) of the ovarian Xenium sample displaying cell boundaries (white lines), bin size overlies (pink squares) and transcript locations for *KRT7* (yellow dot). **Top Right:** Averaged density plot of binning simulation in Xenium samples, showing the number of different cells with transcripts in each bin**. Bottom Right:** Averaged density plot of binning simulation in Xenium samples, showing the number of bins overlapping with transcripts from the same cell. **b)** Transcript count per bin in a small region of the ovarian sample in VisiumHD. **c)** Scatter plots illustrating the correlation between density gene expression in VisiumHD (x-axis) and CytAssist (y-axis, top) or Xenium (y-axis, bottom. The dashed diagonal line indicates expected perfect correlation (x = y). **d) Top:** Box plot comparing the spatial distribution (Moran’s I) of genes in VisiumHD (∼18K genes) and Xenium (377 genes). The dots are colored according to the distance to the diagonal (equal Moran’s I across platforms). **Bottom:** Scatter plot comparing the spatial distribution (Moran’s I) of shared genes in VisiumHD and Xenium (∼370 genes). The dots are colored according to the distance to the diagonal (equal Moran’s I across platforms). **e)** Expression of *CXCL10* (top; Moran’s I: 0.59 in Xenium, 0.36 in VisiumHD) and *EPCAM* (bottom; Moran’s I: 0.68 in Xenium, 0.10 in VisiumHD) in the ovarian sample, with Xenium data shown on the left and VisiumHD on the right **f)** Barplot showing the cell type relative abundance found in the whole VisiumHD sample (left) and the cropped Xenium area (right) in each tumor sample. **g)** Barplot displaying the difference in absolute cell type abundance between the full VisiumHD sample and the cropped Xenium area for each tumor sample. Bars are color-coded by cell type, matching the palette used in f). Positive values indicate higher abundance in VisiumHD, while negative values indicate higher abundance in Xenium. **h)** Cell type characterization of Cancer (epithelial), Stroma cells (fibroblast, smooth muscle and endothelial), T cells, Macrophages and Immune (remaining cell types) in the ovarian sample (left). Localization of two macrophage subpopulations with a pair of overexpressed markers on the top (right). **i)** Score of Hypoxia pathway in the ovarian sample using VISIUM CytAssist (top) and VisiumHD (bottom).

As expected, the choice of binning size can have a significant impact on the data. In terms of cell mixture, the 2 µm bins have, on average, less than one cell per bin, whereas the 8 µm per side have, on average, transcripts from 2 to 4 different cells, and the 16 µm bins between 5 and 10 different cells, though this range varies depending on the tumor type and its composition (**Figure S14**). Regarding the number of bins per cell, at the highest resolution (2 µm), most cells occupy between 10 and 25 bins, whereas at 8 µm and 16 µm these ranges increase to 2-7 and 1-5, respectively. Tools such as Bin2Cell^29^ address this limitation by aggregating 2×2 µm bins into single-cell units using hematoxylin and eosin (H&E) images alongside gene expression data.

The transcriptional data produced by VisiumHD correlated well with tissue morphology. In agreement with the simulated binning experiment with the Xenium data, the range of median transcripts per bin in most samples was similar (∼100 to ∼300), except for the lung sample, which had an order of magnitude less median transcripts **(Figure S15**). That being said, when plotting the number of transcripts per bin, we observed a technical artifact resembling the grid pattern seen in CosMx, but now showing as row and column “striping”. This has been previously noted in the Bin2Cell study, and needs to be taken into account in downstream analyses. We then looked at the correlation between the average gene expression in VisiumHD, and VISIUM CytAssist and Xenium. The overall correlation with both platforms is quite good (R > 0.97 with CytAssist and R > 0.85 with Xenium, **Figure 7c).** However, the UMI density in VisiumHD was ∼3 times lower than VISIUM CytAssist and ∼15 times lower than Xenium. Next, we looked at the spatial autocorrelation of the transcriptional data with the Moran’s I. In this case, we only compared VisiumHD with Xenium, as differences in resolution can bias this metric, and would make the comparison with VISIUM CytAssist not meaningful. Our results show that the Moran’s I in Xenium is significantly higher for all shared genes than in VisiumHD **(Figure 7d**). This can probably be attributed to a higher sparsity of the data, probably caused by the whole-transcriptome coverage. Despite this discrepancy, gene expression data in VisiumHD correlated with morphological structures within tissue **(Figure 7e**). These findings underscore the need to account for the high sparsity of VisiumHD data when interpreting gene expression patterns.

Finally, we looked at the power of VisiumHD data to identify different cell types and explore the expression levels of different pathways, two common tasks in spatial transcriptomics projects. The bin-level cell type classification using SingleR reproduced the tissue morphology seen in Xenium (**Figure S16**). However, we found notable shifts in cell type composition between the two technologies. In VisiumHD, we observed a higher relative abundance of epithelial and stromal cells compared to immune cells (**Figure 7f**). This difference could be due to cell size, as larger cells are more likely to span multiple bins, as reflected in absolute counts (**Figure 7g**). In the case of more granular cell types, such as T and B cells, the proportions were generally comparable across platforms, with some exceptions. For instance, the lymphoma sample lacked effector T cells in VisiumHD, potentially due to reduced capture of the *CCR7* marker (**Figure S17, Figure S18**). Next, we used unsupervised clustering to test if the whole-transcriptome data from VisiumHD allowed us to identify rarer cell types not captured in Xenium. Among others, we were able to identify inflammatory, *SPP1*+ and *SELENOP*+ macrophages in the ovarian sample, which were missed in the Xenium data (**Table S1**). That being said, we also identified several macrophage subpopulations in both platforms that clustered largely by the expression markers of neighboring cells. For example, we observed *EPCAM*+ *SOX17*+ macrophages in Xenium and *WFDC2*+ *MUC16*+ macrophages in VisiumHD, both showing colocalization with cancer cells (**Figure 7h**). This pattern was also seen in other cell types across both platforms. This is likely due to either bin mixture in VisiumHD or the limited transcript coverage and potential misassignment in Xenium.

A key approach in studying tumor systems involves analyzing groups of functionally related genes or pathways. These pathways provide a comprehensive view of the complex molecular interactions that drive tumor behavior, extending beyond simple cell type classifications. For instance, we investigated the Hypoxia pathway in ovarian tissue samples, which is known to influence the TME, promote mesenchymal phenotypes, and contribute chemoresistance through metabolic alterations. Low gene coverage platforms can limit this type of analysis due to insufficient overlap between the gene panel and the pathway gene signature, potentially leading to misleading conclusions. By contrast, platforms with whole-transcriptome coverage allow for more reliable pathway scoring. For example, we quantified this score across VISIUM CytAssist and VisiumHD platforms, revealing a similar landscape of tumor heterogeneity (**Figure 7i**). However, VISIUM CytAssist resolution remains a limiting factor in hypothesizing the associations between cell state, cell identity, and cell neighborhood accurately. In summary, while VisiumHD data has inherent limitations, primarily due to its binning approach, it provides a valuable opportunity to explore cancer tissues from novel spatial perspectives.

### Correlation of spatial transcriptomics and spatial proteomics

While RNA and protein levels for most genes tend to be highly correlated, this is not always the case. This is important, as oftentimes the molecular functions of any given gene are carried out not by its transcript, but by the protein it encodes. Therefore, in those cases where RNA and protein do not correlate, assessing the RNA to infer molecular function can be misleading. This includes important phenomena in cancer, such as disruption of protein-protein interactions by driver mutations, or lowering the levels of tumor suppressors by germline or somatic variants.

To assess to what extent spatial transcriptomics correlates with protein levels, we generated matching spatial proteomics data using the VISIUM CytAssist immuno-oncology panel, which includes 35 proteins. This panel can be generated during the regular VISIUM CytAssist protocol for spatial transcriptomics, and relies on antibodies tagged with a nucleotide sequence which are then labeled with the spatial barcode from the VISIUM spot. In general terms, our five samples showed a weak correlation between gene and protein expression (**Figure 8a**) across all studied genes, with an average correlation coefficient around ∼0.1. This is in stark contrast to the average correlation between RNA and protein levels for these genes in bulk data from the Clinical Proteogenomics Tumor Analysis Consortium across 7 cancer types (**Figure 8b**), where the average correlation coefficient is ∼0.7.

**Figure 8.**
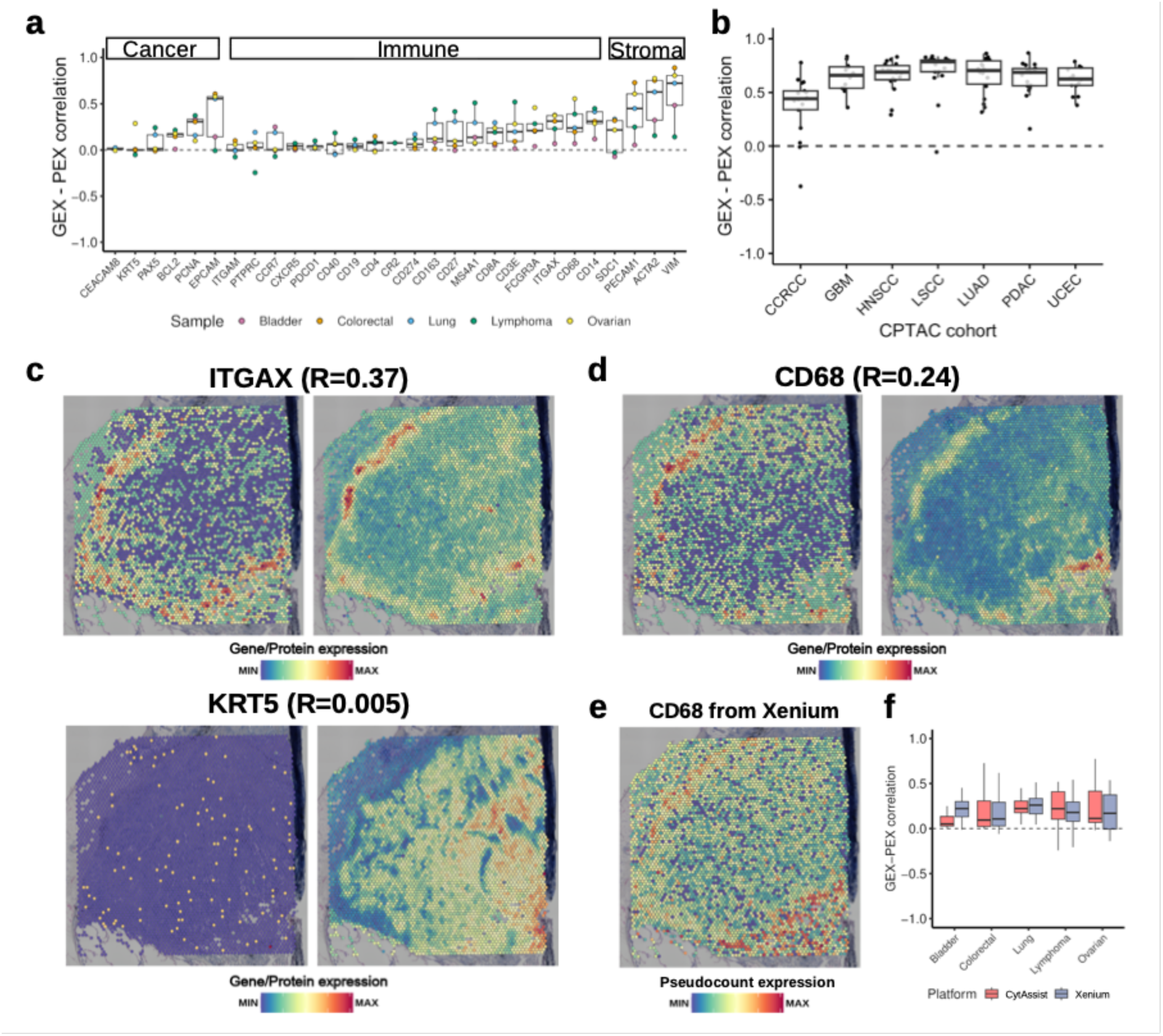
Spatial multi-omics reveals differences in RNA and protein levels across space. **a)** Boxplots showing the correlation between normalized gene and protein expression in each sample represented by a color. Genes are sorted based on which cell type is expected to be overexpressed and their median correlation. **b)** Boxplot showing the correlation between RNA and protein for the 35 genes included in the CytAssist protein panel, according to the bulk measurements in 7 CPTAC cancer types. **c)** Normalized gene (left) and protein expression (right) of *ITGAX* (top) and *KRT5* (bottom) in the Lung sample. Correlations between gene and protein expression are shown alongside gene symbols. **d)** Normalized gene (left) and protein expression (right) of *CD68* in the Lung sample. Correlation between gene and protein expression are shown alongside gene symbol. **e)** Normalized gene expression of *CD68* after transferring Xenium transcripts to CytAssist spots. **f)** Same as b) but correlations are computed using gene expression from CytAssist and the pseudo counts from Xenium. In both cases, only genes found in the protein and Xenium panel are considered.

When considering individual genes, we identified several distinct scenarios. First, positive correlations were observed between genes and proteins in cases where genes were sufficiently captured and exhibited a spatial distribution such as *ITGAX* in the lung sample (**Figure 8c, top**). Conversely, genes with correlations close to zero, predominantly cancer and immune-related genes, tended to have low RNA expression, as illustrated by the example of *KRT5* in the lung sample (**Figure 8c, bottom**). Last, we encountered a third scenario where genes and proteins were both highly captured and had a similar spatial distribution but exhibited regions of overexpression. For instance, *CD68* showed a high gene expression uniquely in the upper-left part of the tissue adjacent to the adipose tissue (**Figure 8d**). While at the protein level, it also expressed an overlap in this area, but additionally showed high expression in the bottom-right area of the sample. Initially, we hypothesized that the differences between hotspots could be attributed to biological processes (i.e. mRNA translation or degradation) or the presence of distinct macrophage populations. However, we compared the gene expression of Xenium by creating pseudo spots after image registration of Xenium and CytAssist samples. Despite observing a slightly noisy spatial pattern, we found that regions with high expression of RNA (as inferred from Xenium) and protein were largely concordant, suggesting some differences in the quantification of the same gene by VISIUM and Xenium (**Figure 8e**). We explored whether this phenomenon extended to other genes, but our results show that, except for the bladder sample, this seems to be relatively rare (**Figure 8f**).

Overall, there are some intriguing and probably biologically relevant dynamics between the RNA and protein of several of the 35 genes we have studied. Such dynamics cannot be seen in bulk data, as evidenced by the CPTAC analysis. Instead, they can only be seen in spatial -omics platforms. Therefore, the future of the field must further the development of assays that allow the capture of multiple -omics readouts.

## Discussion

Spatial -omics technologies, and spatial transcriptomics in particular, are transforming our understanding of many biological phenomena^30^, providing insights into key questions that have important medical applications^31^. While these technologies will virtually impact every field of biological research, one of the most promising applications is in cancer research^32^, where it will help us to decipher the inner workings of tumors, the spatial organization of heterogeneous cancer cells, and how they interact with the tumor microenvironment^33^.

There are now many different spatial transcriptomics platforms available, making it difficult for researchers not yet familiar with the topic to decide which exact platform to use. To that end, here we have made a systematic comparison of five different spatial transcriptomics platforms when applied to FFPE human tumor samples from six cancer types. We focused on FFPE specimens as these are the starting material for most cancer research projects. Furthermore, whenever possible, we sectioned the samples for the different platforms on the same day to limit the impact of batch effects and maximize the comparison of the histological sections.

We first would like to acknowledge the limitations of our study. First and foremost, the field of spatial transcriptomics (and spatial biology in general) is advancing at an incredible speed, so the results presented here should be interpreted in the context of the state of the field in early 2024. For example, 10x Genomics now provides membrane staining for Xenium, an option not available at that time and that can improve cell segmentation. Similarly, both CosMx and Xenium now allow users to run panels with significantly higher plex than the ones available to us at the moment we did our experiments, including 5000 genes in the case of Xenium and 6000 for CosMx (with a scheduled release of the whole-transcriptome panel by 2025). Finally, we did not include in our comparison a variety of other spatial transcriptomics platforms.

It is also important to note that there have been several recent studies comparing spatial transcriptomics platforms. For example, Cook *et al.*, compared four serial sections of an FFPE prostate cancer sample, two of them analyzed with Xenium and the other two with CosMx^34^. Similarly, Wang *et al.* compared three *in situ* platforms (CosMx, Xenium, and MERSCOPE) across 23 different tissues using serial sections from a tissue microarray containing both, normal as well as tumor tissue samples^35^. More recently, Hartman and Satija compared six *in situ* platforms using data from fresh frozen mice brain samples^36^, and You *et al* compared 11 sequencing-based spatial transcriptomics platforms with a focus on mice samples^37^.

None of these previous analyses included matching VISIUM data in either of its versions (manual, CytAssist, or HD), so we can only compare our results regarding the benchmarking of *in situ* platforms (CosMx and Xenium). In that regard, our findings largely align with those previously described. In brief, Xenium does seem to have a lower false discovery rate than CosMx, particularly in lowly expressed genes. We also showed that this affects the actual spatial distribution of the signal of many genes, as reflected by their Moran’s I, which tends to be lower in CosMx than in Xenium. Also agreeing with previous studies, the segmentation data from CosMx seems to yield smaller cells with sizes more similar to those observed in actual cells, probably thanks to the membrane staining. Nevertheless, when looking at cell typing, these differences in false discovery rate and segmentation do not seem to affect the overall results, as both platforms estimate similar cell fractions in most samples.

There are several aspects, though, that are unique to our study. Among others, we have done the first systematic comparison between the manual and the CytAssist versions of VISIUM. Our results show that there is, indeed, a significant improvement in overall data quality when using the CytAssist instrument, as evidenced by the higher number of detected genes as well as the higher spatial autocorrelation of their associated transcripts. This can be attributed to both, better overall chemistry of the new version (with three probes per gene instead of one), as well as a potential improvement in analyte transfer thanks to the CytAssist. This can be seen in Figure 6c, where the area of cells that, according to Xenium, are expressing *ACTA2* but are located within spots, are not detected at all with VISIUM.

Another novel aspect of our study is the quantification of the impact of tissue area analyzed in establishing the cell type composition of a sample. This is important as, to optimize precious resources (such as clinical samples), many studies using spatial transcriptomics tend to analyze a relatively small area of each sample to, instead, increase the number of samples. One typical example is the use of tissue or tumor microarrays, where one only studies a circle between 0.6mm and 1.2mm in diameter, which is significantly smaller than the capture area of VISIUM (6.5mm x 6.5mm), but allows fit more than one sample in each space, increasing the total number of analyzed samples. In line with previous studies working on spatial proteomics, we showed that reducing the analyzed area can lead to extremely biased cell composition estimates. We recommend maximizing this number to get a better representation of the tissue.

To the best of our knowledge, this is also the first study to report a comparison of spatial transcriptomics platforms including VisiumHD data for serial sections of the same samples. This platform is currently the only one that can deliver whole-transcriptome coverage at single-cell resolution. Our results show that the data is of very good quality, effectively delivering both promised features. Whoever, it does have some challenges. On the one hand, despite the fact that we sequenced above the minimum recommended reads per spot, we did not reach saturation of the libraries, and this probably was reflected in the sparsity of the expression patterns of many genes as well as the significantly lower number of UMIs per area than in Xenium and VISIUM CytAssist. That being said, the overall structure of the expression patterns was very similar to that obtained with the other single-cell resolution platforms (Xenium and CosMx), while having whole-transcriptome coverage. Furthermore, the overall estimated cell composition was also similar. The other significant challenge with data from VisiumHD is the assignment of the transcripts of each bin to the correct cell. While this is shared with virtually any other platform working at single-cell resolution, it becomes evident when looking at different clusters of the same cell type. Some of these clusters were driven by expression of markers from other cell types neighboring the ones that we were studying. To extract all the power from this platform, the regular grid-binning provided by 10x will have to be changed with other tools that leverage the H&E segmentation to properly assign each bin to the right cell.

Finally, by using the VISIUM proteomics panel, we were able to compare the spatial distribution of transcripts and their corresponding proteins for the same tissue sample, revealing some very interesting dynamics. While in some cases we did find that both, protein and transcript, correlate in space, these were only a minority. In several cases (such as *KRT5*) there was no correlation whatsoever between both. These were frequently caused by the lack of detection of RNA despite the presence of protein in the tissue. In other cases, the correlation between RNA and protein differed depending on the specific location within the sample, suggesting that the transcription / translation dynamics of these genes might differ depending on the cell type. Given the relevance of protein assays in the clinical setting, particularly the widespread use of immunohistochemistry in pathology laboratories for different diagnostic procedures, these results show the importance of spatial multi-omics characterization of tumor samples, as it is the only way to identify these different scenarios.

Overall, as additional technologies for spatial transcriptomics become available, it will be essential to perform rigorous comparisons to help the community prioritize each of them based on their strengths, limitations, and the ultimate aims of the studies.

## Acknowledgments

We thank the patients who generously donated their samples. We also would like to thank Luciano Martelotto for helpful discussions of the results. This work was supported by Nanostring and Longwood Laboratories (who funded the CosMx experiments), 10x Genomics and Bonsai Labs (who funded the Xenium experiments), and Asociación Española Contra el Cáncer (who funded the VISIUM experiments, LABAE20038PORT). S.C is supported by AECC (LABAE20038PORT). E.P-P is supported by the Spanish Ministry of Science (RYC2019-026415-I), a Fundación FERO fellowship (BFERO2022.6), and Departament de Recerca i Universitats / Generalitat de Catalunya (2021 SGR 01309).

## Author contributions

Study conception and design: E.P-P, D.G, M.E; performed experiments and data collection: D.G, E.P., E.M., F.R., S.C; computational analyses: S.C, E.P-P; writing, review, and editing: S.C, M.E, E.M., F.R., D.G, E.P-P.

## Declaration of interests

The authors declare no competing interests beyond the company-sponsored experiments disclaimed in the acknowledgments section.

## Methods

### Sample acquisition

All tumor samples where obtained under the ethics committee approval 2022/78-APA-HUGC

### Tissue preparation and sequential sectioning

For our benchmarking purpose, we selected six untreated primary tumor samples, covering the most common cancer types: colorectal, breast, lung, lymphoma, ovarian, and bladder. The H&E staining from each formalin-fixed paraffin-embedded (FFPE) blocks, helped us to characterize the tissues and confirmed that each sample had an adequate and good quality representation, not only of tumor tissue but also of the surrounding healthy tissue, to warrant the experiments.

First, the quality of preserved RNA in the FFPE blocks was evaluated based on the percentage of RNA fragments above 200 base pairs (DV200). For each block, we sectioned and collected three consecutive 8 μm sections for RNA extraction using the RNeasy FFPE kit for RNA Purification (Qiagen Cat No: 73504). We analyzed the purified RNA with Agilent tapestation, and chose the samples with values above 34% of DV200. The dimensions of the tissue in the selected blocks were approximately 7 mm by 7 mm, except in the bladder sample that the block was formed by small pieces of tissue from the same patient.

To facilitate a comprehensive comparison of data from VISIUM manual, Xenium, CosMx, and VISIUM CytAssist platforms, we sectioned the chosen samples as illustrated in Figure 1b. More in detail, from each of these samples we had collected and processed one 7um section for VISIUM manual analysis 6 months ago, and we had stored the blocks at 4 degrees. For the benchmarking study we retrieved the blocks, discarded between 5 to 10 - 5um sections and collected three 7 μm serial sections for Xenium, CosMx, VISIUM CytAssist analysis and one extra for H&E staining.

The sectioning was conducted using a semi-automated microtome (ThermoScientific HM340E), utilizing a fresh blade and ensuring thorough decontamination with RNAse AWAY (ThermoFisher Scientific). We detail the sectioning and subsequent slide preparation methods for Xenium, CosMx and VISIUM Cytassist platforms in each chapter.

### Xenium sample preparation

The Xenium workflow began by sectioning 7 μm FFPE tissue sections onto a Xenium slide, according to the “Xenium In Situ for FFPE-Tissue Preparation Guide” (CG000578 Rev C, 10X Genomics). Briefly, the sections were cut, floated in an RNAse-free water bath at 42 degrees, and carefully placed onto the capture area of a Xenium slide (PN-1000465). We strategically placed three 7x 7mm sections in each slide. After sectioning, the slides were incubated at 42 degrees per 3 hours, and kept in a sealed bag with desiccators at 4 degrees overnight. The next day, the slides were shipped to the Dresden Genome Center Facility, where the experiments were conducted.

At the Dresden Genome Center Facility, the Xenium slides were processed following the “Xenium In Situ for FFPE-Deparaffinization and Decrosslinking” protocol (CG000580 Rev C, 10X Genomics). The slides were equilibrated to room temperature and the protocol of deparaffinization was performed.

Subsequently, the Xenium slides were assembled into Xenium cassettes (PN-1000566, 10X Genomics), which allow for the incubation of slides on the Xenium Thermocycler Adapter inside a Thermocycler machine with a closed lid for optimal temperature control. The slides were processed using the “Xenium Slides and Sample Prep Reagents” kit (PN-1000460, 10X Genomics), starting with incubation in a decrosslinking and permeabilization solution at 80 °C for 30 minutes, followed by a wash with PBS-T.

The Xenium slides were then processed according to the “Xenium In Situ Gene Expression” user guide (CG000582 Rev D, 10X Genomics). The slides were incubated at 50 °C overnight for aproximately 19 hours with the gene expression panel (“Xenium Human Multi-Tissue and Cancer Panel”, PN-1000626, 10X Genomics, which targets 377 human genes). This was followed by a series of washes and steps, including a post-hybridization wash at 37 °C for 30 minutes, a ligation at 37 °C for 2 hours, and an amplification step at 30 °C for 2 hours. After additional washing steps, the slides were treated with an autofluorescence quencher and a nuclei staining step.

Finally, at the end of the second day of the protocol, the two slides in the cassettes were loaded into the Xenium Analyzer. The first step in the instrument consists in a sample scan, were images of the fluorescent nuclei in each section are given, and these images allow the user to select and determine the regions to be included in the analysis. For the 6 samples, we selected all the tissue to be analyzed.

The run in the Xenium Analyzer lasted around 50 hours. After that, the instrument was emptied of consumables and the Xenium slides carefully removed. PBS-T was added to the slides and a post-run H&E staining was performed.

### Visium CytAssist whole-transcriptome and protein panel

Visium CytAssist with the protein panel provides whole-transcriptome, spatially-barcoded sequence data. The histology workflow was performed using the Visium CytAssist Spatial Gene Expression for FFPE (Demonstrated Protocol CG000520, 10x Genomics). The tissue was sectioned as described in Visium CytAssist Spatial Gene Expression for FFPE – Tissue Preparation Guide (Demonstrated Protocol CG000518, 10x Genomics). 7 μm sections were cut, floated in an RNAse-free water bath at 42 degrees, and carefully placed on a Microscope Slide (Epredia, cat # 10149870) the slides were incubated at 42 degrees per 3 hours and kept in a sealed bag with desiccators at room temperature overnight.

On the day of the experiment, the Microscope slides containing tissue sections were deparaffinized and stained for H&E. Samples were imaged using a Nikon microscope (Nikon Eclipse Ti2), after imaging with brightfield, the coverslip was removed, followed by hematoxylin destaining and decrosslinking (Demonstrated Protocol CG000520, 10x Genomics). Afterwards, Microscope slides with tissue sections were incubated with whole transcriptome human probes overnight for hybridization, then probe ligation was performed, and probe release was enable using Visium CytAssist instrument to transfer analytes from tissue to a Visium CytAssist Spatial Gene Expression slide with 6.5 x6.5 mm capture areas.

The library construction for gene and protein expression was done following 10x genomics User Guide CG000494. Briefly after probe ligation, samples were incubated with the protein panel, and two libraries were generated per sample.

Gene Expression and Protein Expression Libraries were sequenced with paired-end dual-indexing (28 cycles Read 1, 10 cycles i7, 10 cycles i5, 50 cycles Read 2). Sequencing libraries were demultiplexed with bcl2fastq (Illumina). The Space Ranger pipeline v2.1 (10x Genomics) and the GRCh38-2020-A reference were used to process FASTQ files.

### CosMx data generation

The FFPE blocks from each sample were sectioned as described in CosMx Tissue Preparation Guide (MAN-10184-02, Nanostring). Briefly, 7 μm sections were cut, floated in an RNAse-free water bath at 42 degrees, and carefully placed onto the center of a Microscope Slide (Superfrost Plus, Epredia J1800AMNZ), considering the specific area the CosMx instrument can analyze. After sectioning, the slides were incubated at 42 degrees per 3 hours, and kept in a sealed bag with desiccators overnight. The slides were shipped to NanoString Seattle for profiling as part of their early technology access program (TAP).

Once in Seattle, the slides were processed following Nanostring MAN-10184-02. Briefly, the samples were baked overnight at 60 °C, followed by preparation for in-situ hybridization (ISH) on the Leica Bond RXm system by deparaffinization and heat-induced epitope retrieval (HIER) at 100 °C for 15 min using ER2 epitope retrieval buffer (Leica Biosystems product, Tris/EDTA-based, pH 9.0). After HIER, tissue sections were digested with 3 μg/ml Proteinase K diluted in ACD Protease Plus at 40 °C for 30 min.

Tissue sections were then washed twice with diethyl pyrocarbonate (DEPC)-treated water (DEPC H2O) and incubated in 0.00075% fiducials (Bangs Laboratory) in 2X saline sodium citrate, 0.001% Tween-20 (SSCT solution) for 5 min at room temperature in the dark. Tissue sections were fixed with 10% neutral buffered formalin (NBF) for 5 min at room temperature. Fixed samples were rinsed twice with Tris-glycine buffer (0.1 M glycine, 0.1 M Tris-base in DEPC H2O) and once with 1X PBS for 5 min each before blocking with 100 mM N-succinimidyl (acetylthio) acetate (NHS-acetate, Thermo Fisher Scientific) in NHS-acetate buffer (0.1 M NaP, 0.1% Tween pH 8 in DEPC H2O) for 15 min at room temperature. The sections were then rinsed with 2X saline sodium citrate (SSC) for 5 min and an Adhesive SecureSeal Hybridization Chamber (Grace Bio-Labs) was placed over the tissue.

NanoString ISH probes were prepared by incubation at 95 °C for 2 min and placed on ice, and the ISH probe mix (1 nM 980 plex ISH probe, 10 nM Attenuation probes, 1 nM SMI-0006 custom, 1X Buffer R, 0.1 U/μL SUPERase•InTM [Thermo Fisher Scientific] in DEPC H2O) was pipetted into the hybridization chamber and hybridization was performed at 37 °C overnight. Tissue sections were washed twice in 50% formamide (VWR) in 2X SSC at 37 °C for 25 min, washed twice with 2X SSC for 2 min at room temperature, and blocked with 100 mM NHS-acetate in the dark for 15 min. In preparation for loading onto the CosMx SMI instrument, a custom-made flow cell was affixed to the slide.

RNA target readout on the CosMx SMI instrument was performed. Briefly, the assembled flow cell was loaded onto the instrument and Reporter Wash Buffer was flowed to remove air bubbles. A preview scan of the entire flow cell was taken, and 40-58 fields of view (FOVs) were placed on the tissue to match regions of interest. RNA readout began by flowing 100 μl of Reporter Pool 1 into the flow cell and incubation for 15 min. Reporter Wash Buffer (1 mL) was flowed to wash unbound reporter probes, and Imaging Buffer was added to the flow cell for imaging. Nine Z-stack images (0.8 μm step size) for each FOV were acquired, and photocleavable linkers on the fluorophores of the reporter probes were released by UV illumination and washed with Strip Wash buffer. The fluidic and imaging procedure was repeated for the 16 reporter pools, and the 16 rounds of reporter hybridization-imaging were repeated multiple times to increase RNA detection sensitivity.

### Visium HD data generation

Visium HD is a single cell scale resolution whole transcriptome probe-based gene expression assay. The Visium HD workflow began by sectioning 7 μm FFPE tissue sections onto a Microscope Slide (Epredia, cat # 10149870) according to the “Visium HD FFPE Tissue Preparation Handbook” (CG000684 Rev A, 10X Genomics). Briefly, the sections were cut, floated in an RNAse-free water bath at 42 degrees, and carefully placed onto the center of the superfrost slide. After sectioning, the slides were incubated at 42 degrees per 3 hours and kept in a sealed bag with desiccators at room temperature until the day of the experiment.

The slides were then deparaffinized and stained for H&E. The imaging was done using a Nikon microscope (Nikon Eclipse Ti2), after imaging with brightfield, the coverslip was removed, followed by hematoxylin destaining and decrosslinking (Demonstrated Protocol CG000520, 10x Genomics). Afterwards, Microscope slides with tissue sections were incubated with whole transcriptome human probes overnight for hybridization, then probe ligation was performed, and probe release was enable using Visium CytAssist instrument to transfer analytes from tissue to a Visium HD Spatial Gene Expression slide, with 6.5 x6.5 mm capture areas according to User Guide “Visium HD Spatial Gene Expression Reagent Kit” (CG000685 Rev B, 10X Genomics). After probe elution was completed, an amplification step was performed and library preparation was finalized.

After library construction for gene expression was done, libraries were sent for sequencing to Macrogen Korea, and were sequenced with NovaSeq X 10B sequencing (150PE) (43 cycles Read 1, 10 cycles i7, 10 cycles i5, 50 cycles Read 2). Sequenced libraries were demultiplexed with bcl2fastq (Illumina). The Space Ranger pipeline v3.0 (10x Genomics) and the GRCh38-2020-A reference were used to process FASTQ files.

### Data preprocessing of sequencing-based spatial platforms

For VISIUM manual and CytAssist, transcript count matrices from SpaceRanger were imported with Seurat^38^ (v5). Spots were filtered based on their location since there were spots detected out of the tissue or in folded areas with misleading gene expression. Furthermore, genes with less than five transcripts in the whole tissue were discarded. Given that for the breast sample we used the 11.5 mm capture area, we subset the tissue for a fairer comparison with the manual VISIUM. Then, the gene expression was normalized using the SCTransform function from Seurat. Similar preprocessing was done in VisiumHD using 8×8 µm bins resolution. In this case, bins were filtered based on their location and low quantity of transcripts captured.

For CytAssist samples with a protein panel, the protein expression was imported into a new assay independently of the gene expression. No additional spots or proteins were filtered out. Finally, the protein expression was normalized using the centered log ratio (CLR) method implemented in Seurat.

### Data preprocessing of image-based spatial platforms

Gene expression profiles matrices from Xenium Analyzer (v1.5.0.3) of each sample were imported using Seurat (v5). Since the CosMx run aggregated all samples within a single TileDB object, we proceeded to segregate and import each individual sample into a Seurat object based on their respective slide coordinates. During the preprocessing, only cells with no transcripts were filtered. In addition, cells of CosMx that were found in the “Barrel effect” area (see Barrel effect section) were excluded for all comparative experiments. Finally, gene expression was normalized using the SCTransform function from Seurat.

### Image registration across platforms

Since we did not have a whole tissue image for CosMx, we could not apply Image Registration techniques to transfer the coordinates of Xenium to the CosMx. Therefore, we aligned the images by first applying a rotation angle to the Xenium coordinates that was defined by overlaying the distribution of centroids in the space in an image editor software. In the case of the bladder sample, we had to separate the tissue regions as they could differ in orientation and distance. Then we shifted the x and y coordinates of CosMx (since they covered less tissue area) based on common landmarks defined by tissue morphology. Angles and shifting specific for each sample can be found in the code. After that, from the Xenium samples were subset by the limits of each FOV in the CosMx as well we discarded CosMx cells that were not found in the Xenium samples.

In the case of the alignment between CytAssist and Xenium/CosMx, we applied Image Registration using the software Voltron^39^ by defining landmarks between Xenium (DAPI) and CytAssist (H&E) images. Then the transformation was applied in the CytAssist spot coordinates to use them in the Xenium space, so we could transfer the FOV limits and subset.

### Barrel effect

Cells that suffered from the “Barrel effect” in CosMx are found in 36px (4.33 um) from the edge of the FOV. To compare the gene expression in these cells with the rest of the FOV, we subsetted those cells whose centroid coordinates (x or y) concerning the FOV is smaller than 36 or bigger than 4220. Furthermore, to ensure that the differences that we found could be related to the subsetting, we selected an area similar to the barrel effect in the middle of the FOV (between 2000 and 2100 px).

### False Discovery Rate

False Discovery Rate (FDR) in each platform and sample was computed using the following formula:

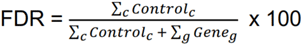

where Control is the sum of the negative control c and Gene is the gene expression of gene g.

### Spatial Autocorrelation

To quantify the spatial variation of the normalized transcripts we use global Moran’s I metric. In VISIUM manual and CytAssist, we used the function CorSpatialFeature from semla package^40^ considering the six closest spots as neighbors (default). For image-based platforms and VisiumHD, the function RunMoransI from the package Voyager_38_. In this case the spatial network was built using the five closest neighbors considering centroid distances.

FindSpatialNeighbors(method=“knearneigh”, dist_type=“idw”, k=5, style=“W”).

### Cell annotation

To annotate image-based sequencing data without reference data, we use SingleR package to map the annotations of the following cell types: epithelial, endothelial, fibroblast, smooth muscle, macrophage, monocyte, dendritic cell, neutrophil, NK cell, B cell, immature B cell, memory B cell, naive B cell, plasma B cell, CD4+ T cell, CD8+ T cell, effector T cell and Treg. The “HumanPrimaryCellAtlasData” was used as a reference and the normalized data with SCT was used as input. Cells with labels NA after the pruning were discarded in the comparison of the technologies.

### FOV sampling experiment

To evaluate the influence of cell type composition within a selected portion of the tissue, we conducted a sampling experiment. Initially, we treated the CosMx FOV size (0.5 mm x 0.5 mm) in the Xenium (shared area) sample as a single unit to facilitate stochastic sampling while adhering spatial constraints. Additionally, we sampled the same number of cells that appear on average in a FOV simulating a single-cell experiment without spatial constraints. Subsequently, we performed 100 iterations of sampling across various quantities of FOVs, including 1, 5, 10, 20, 40, and the maximum number of FOVs possible within the given sample. For each sampling iteration, we calculated the relative abundance of three grouped cell types: cancer (epithelial cells), stroma cells (fibroblasts, smooth muscle and endothelial cells), and immune cells (the remaining cell types).

### Transferring Xenium information to CytAssist spot resolution

After the image registration between Xenium and CytAssist samples, we transfer the coordinates of CytAssist spots to the Xenium space. Then, for each spot, we assigned the cells within a radius of 27.5 μm to the spot center. Finally, for the deconvolution we computed the relative abundance of each cell type from the assigned cells. For the gene expression, we aggregated the sum of gene transcripts found within each spot and we normalized using SCTransform.

### Unsupervised clustering of macrophages

To find macrophages subpopulations in VisiumHD and Xenium, this cell type was subset and renormalized using SCTtransform. Then, dimensional reduction was applied using PCA on the variable genes. Finally cell clusters were identified using FindNeighbors and FindClusters from Seurat using the first 20 PCA dimensions. The resolution used in VisumHD was 0.3 and 0.5 for Xenium to obtain a similar number of subclusters.

### Hypoxia pathway scoring

The transcriptional footprint of the Hypoxia pathway was extracted with PROGENy^41^ using the top 1,000 genes. Then, the pathway activity was inferred using a multivariate linear model implemented in decoupleR^42^ using the normalized gene expression data from VISIUM CytAssist and VisiumHD. For visiualization purposes, q1 and q99 cutoffs of the score were used.

### CPTAC protein and RNA analyses

We downloaded the CPTAC protein and RNA expression matrixes from Liang *et al*^43^.Then, for each gene included in the 10x Genomics CytAssist protein panel, we calculated the correlation coefficient between the bulk RNA counts and the mass-spectrometry protein expression data across all 10 cancer types included in the CPTAC study using a linear model.

## Data availability

All tumor samples where obtained under the ethics committee approval 2022/78-APA-HUGC Raw data can be find at https://nextcloud_admin.carrerasresearch.org/index.php/s/pJzKz4HBE4LPms6/authenticate/showShare with the password 5HG8HtP5.

## Code availability

Code to reproduce the analysis and figures is available at: https://github.com/scervilla/SpatialBenchmarking.

## Notes

### Summary of Updates

We have added additional data for one more spatial transcriptomics platform: Visium-HD

