## Supplementary Figures for "Benchmarking of spatial transcriptomics platforms across six cancer types"

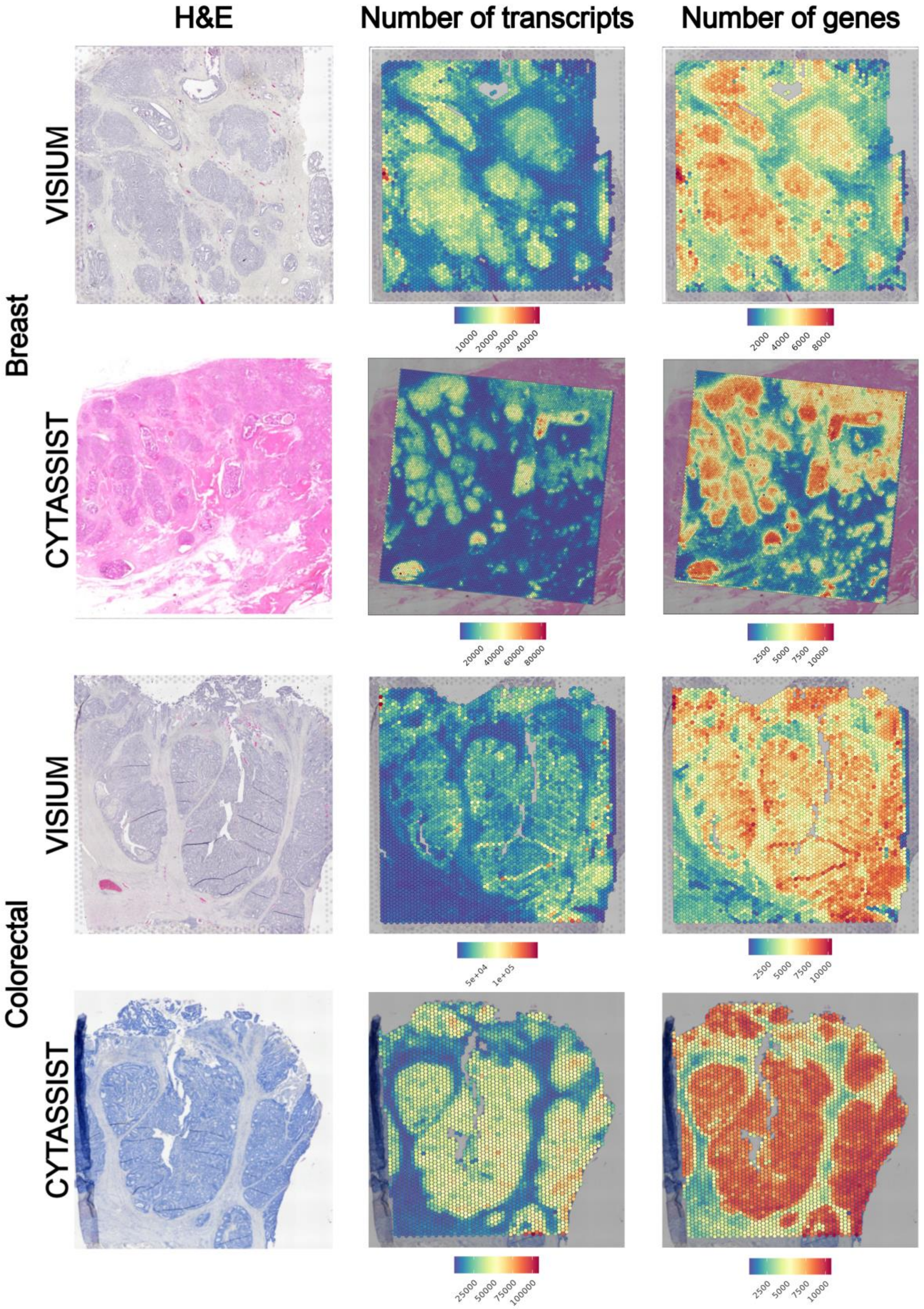
H&E
Number of transcripts
Number of genes
Breast
VISIUM
10000 20000 30000 40000
2000 4000 6000 8000
CYTASSIST
20000 40000 60000 80000
2500 5000 7500 10000
Colorectal
VISIUM
5e+04 1e+05
2500 5000 7500 10000
CYTASSIST
25000 50000 75000 100000
2500 5000 7500 10000

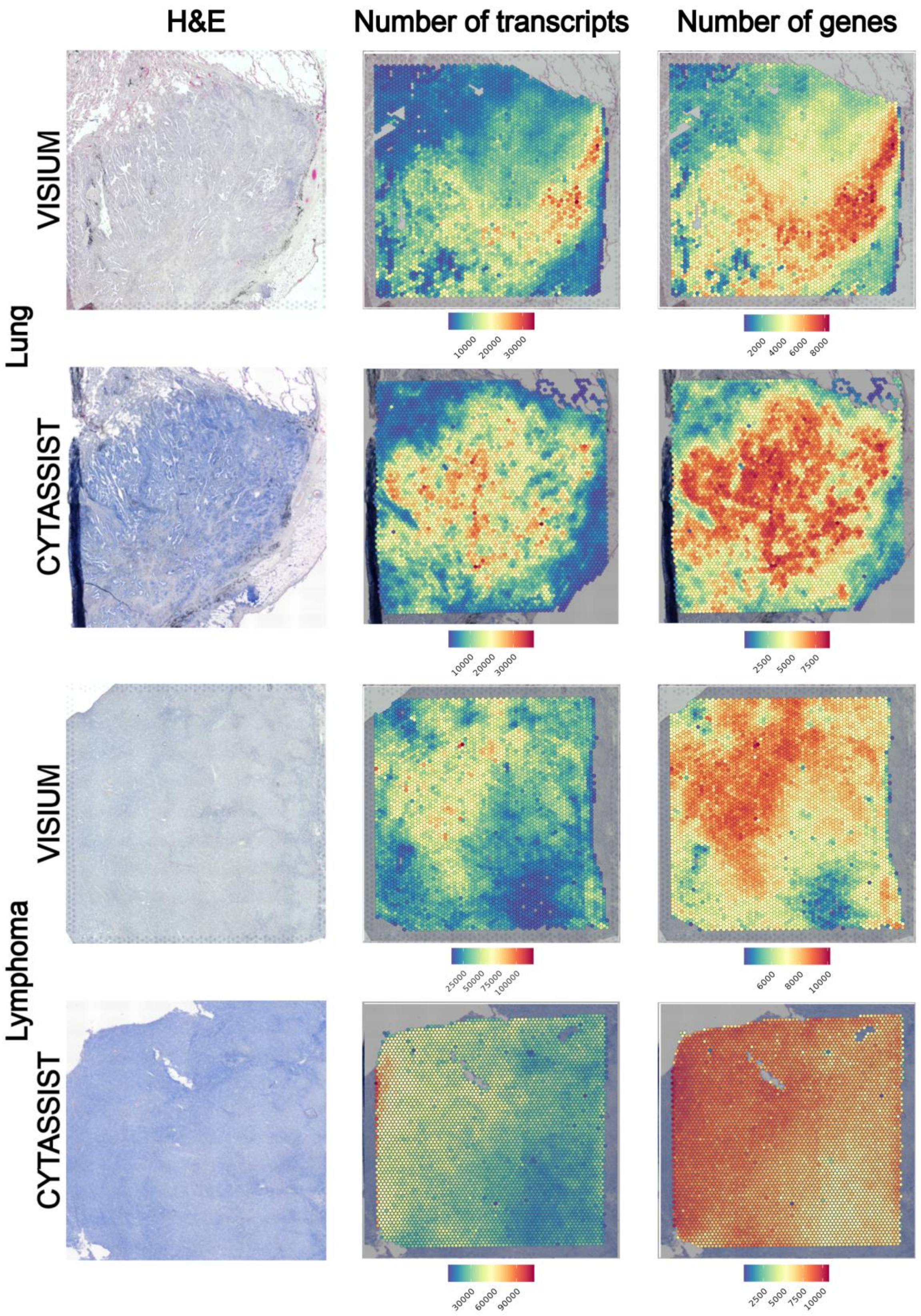
H&E
Number of transcripts
Number of genes
Lung
VISIUM
CYTASSIST
Lymphoma
VISIUM
CYTASSIST
10000
20000
30000
2000
4000
6000
8000
10000
20000
30000
2500
5000
7500
25000
50000
75000
100000
6000
8000
10000
30000
60000
90000
2500
5000
7500
10000

**Figure S1:** Quality Control of Visium manual (top) and CytAssist (bottom)**. Left:** H&E. **Center:** Number of unique genes per spot. **Right:** Number of transcripts per spot.

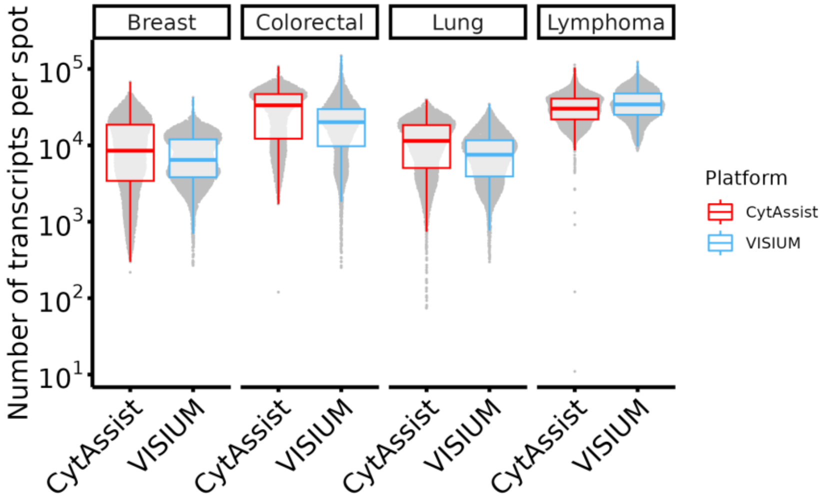


**Figure S2:** Distribution of the number of transcripts detected in each spot across samples and platforms.

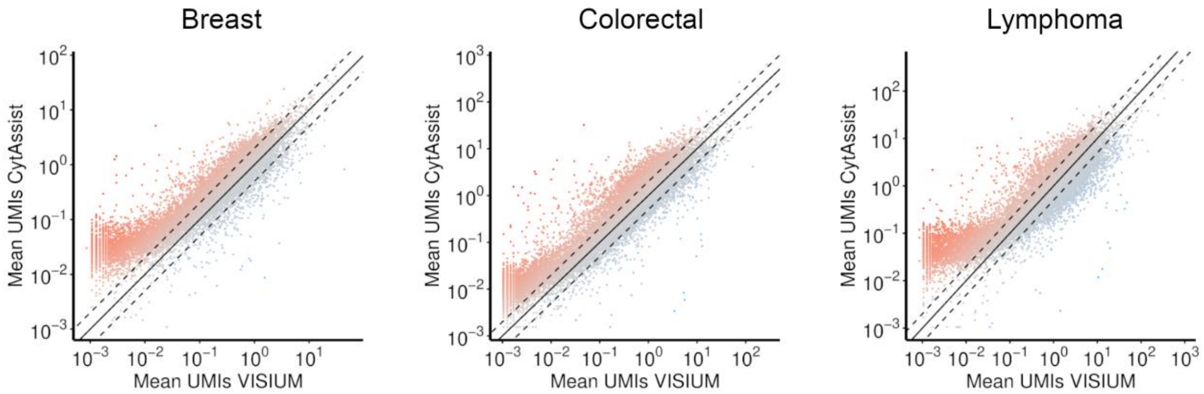


**Figure S3:** Scatter plot of median UMI per genes in VISIUM and CytAssist in the Breast, Colorectal and Lymphoma samples. Color intensity of dots represents the distance from the diagonal (equal UMI detection across platforms).

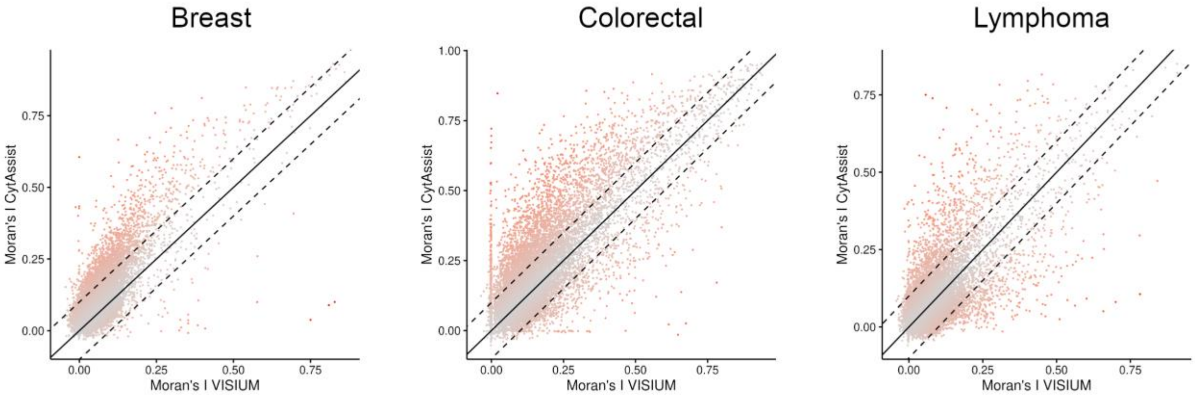


**Figure S4:** Scatter plot of global Moran's I for each gene in VISIUM and CytAssist in the Breast, Colorectal and Lymphoma samples. Color intensity of dots represents the distance from the diagonal (equal UMI detection across platforms).

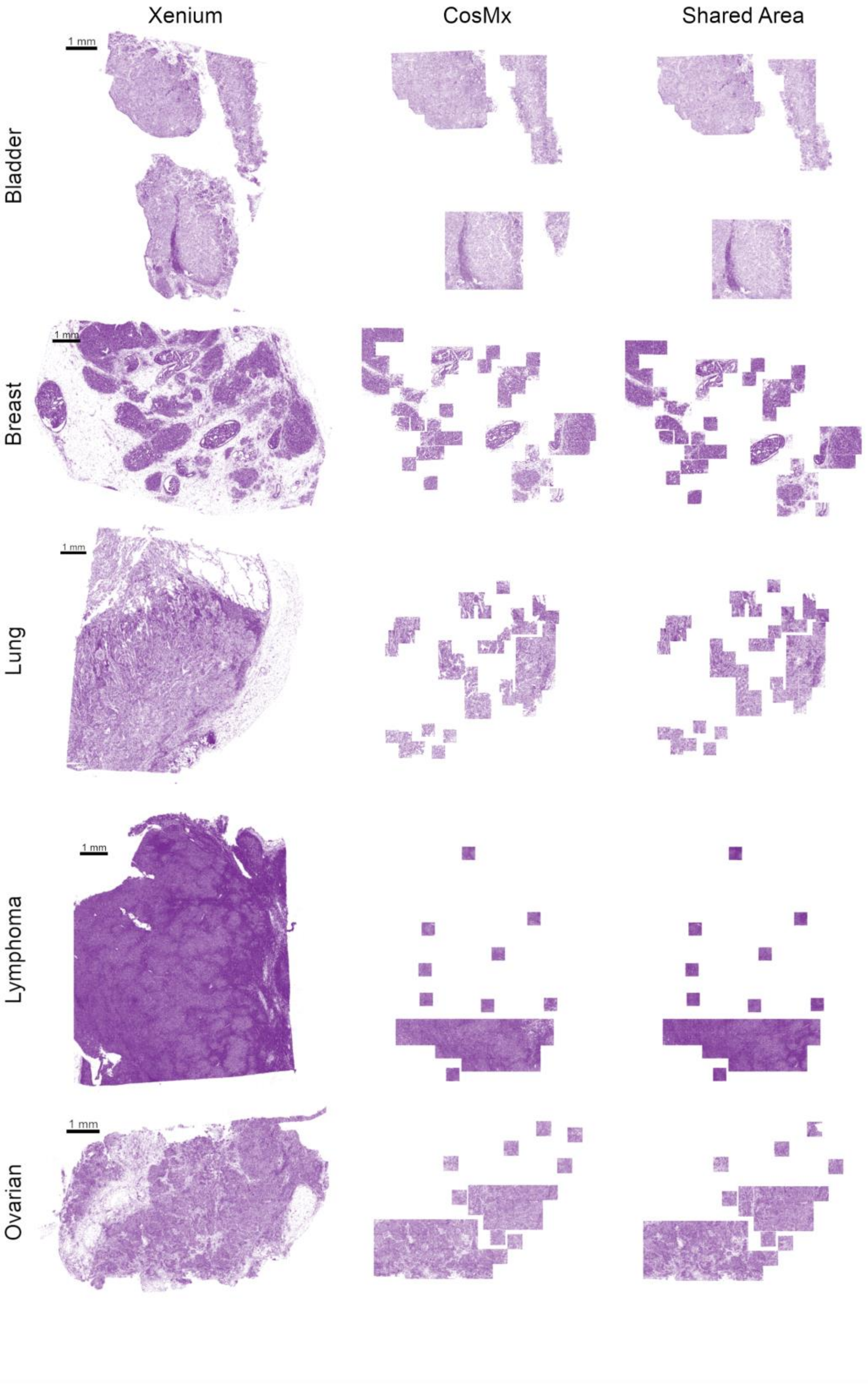


**Figure S5:** Representation of the tissue analyzed and compared in different samples: cells captured using Xenium (left), cells captured using CosMx (middle) and cells that are in the shared regions between Xenium and CosMx (right).

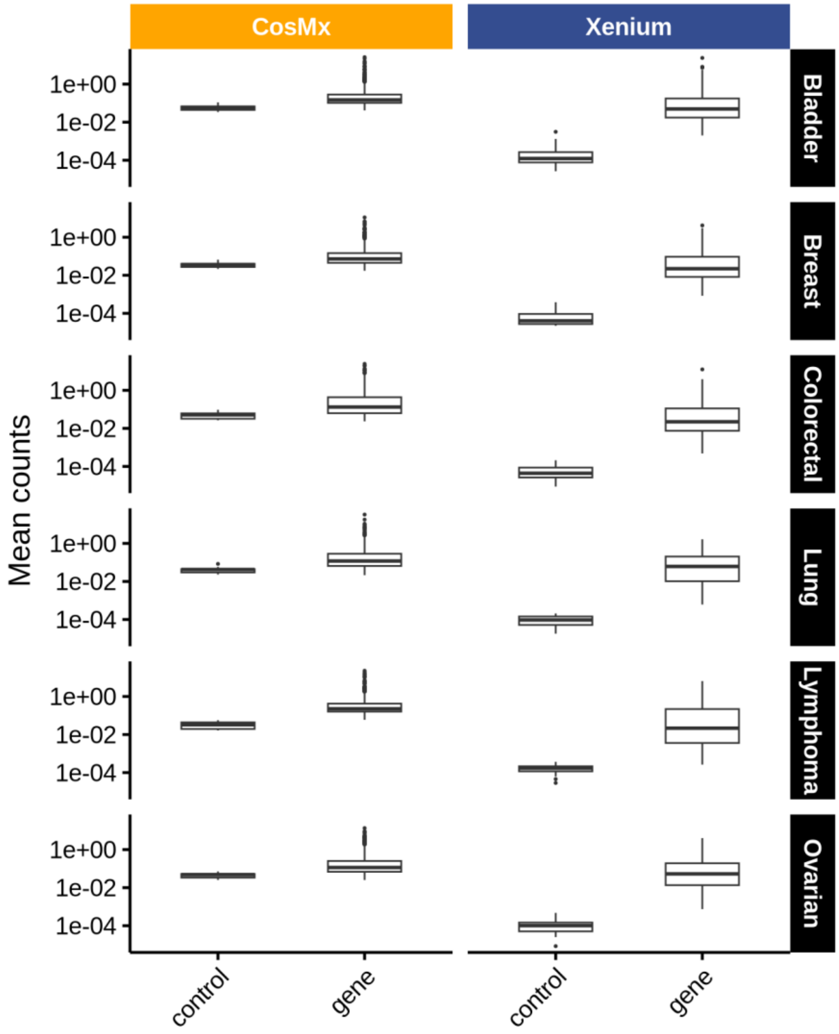


**Figure S6:** Boxplots showing the mean expression of each negative control probe and each gene in CosMx and Xenium.

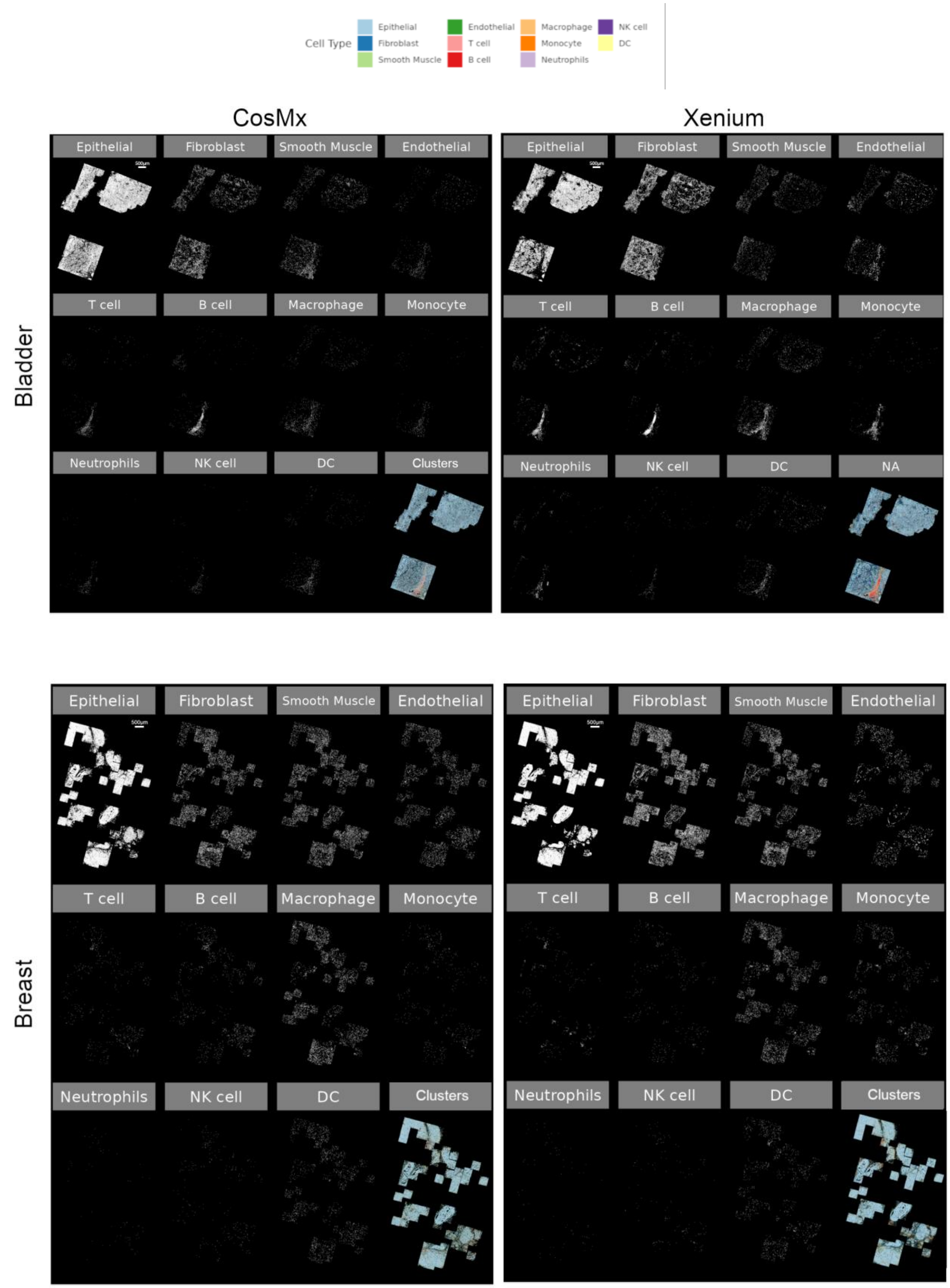

Cell Type
Epithelial
Fibroblast
Smooth Muscle
Endothelial
T cell
B cell
Macrophage
Monocyte
Neutrophils
NK cell
DC
CosMx
Xenium
Bladder
Breast
Epithelial
Fibroblast
Smooth Muscle
Endothelial
T cell
B cell
Macrophage
Monocyte
Neutrophils
NK cell
DC
Clusters
NA

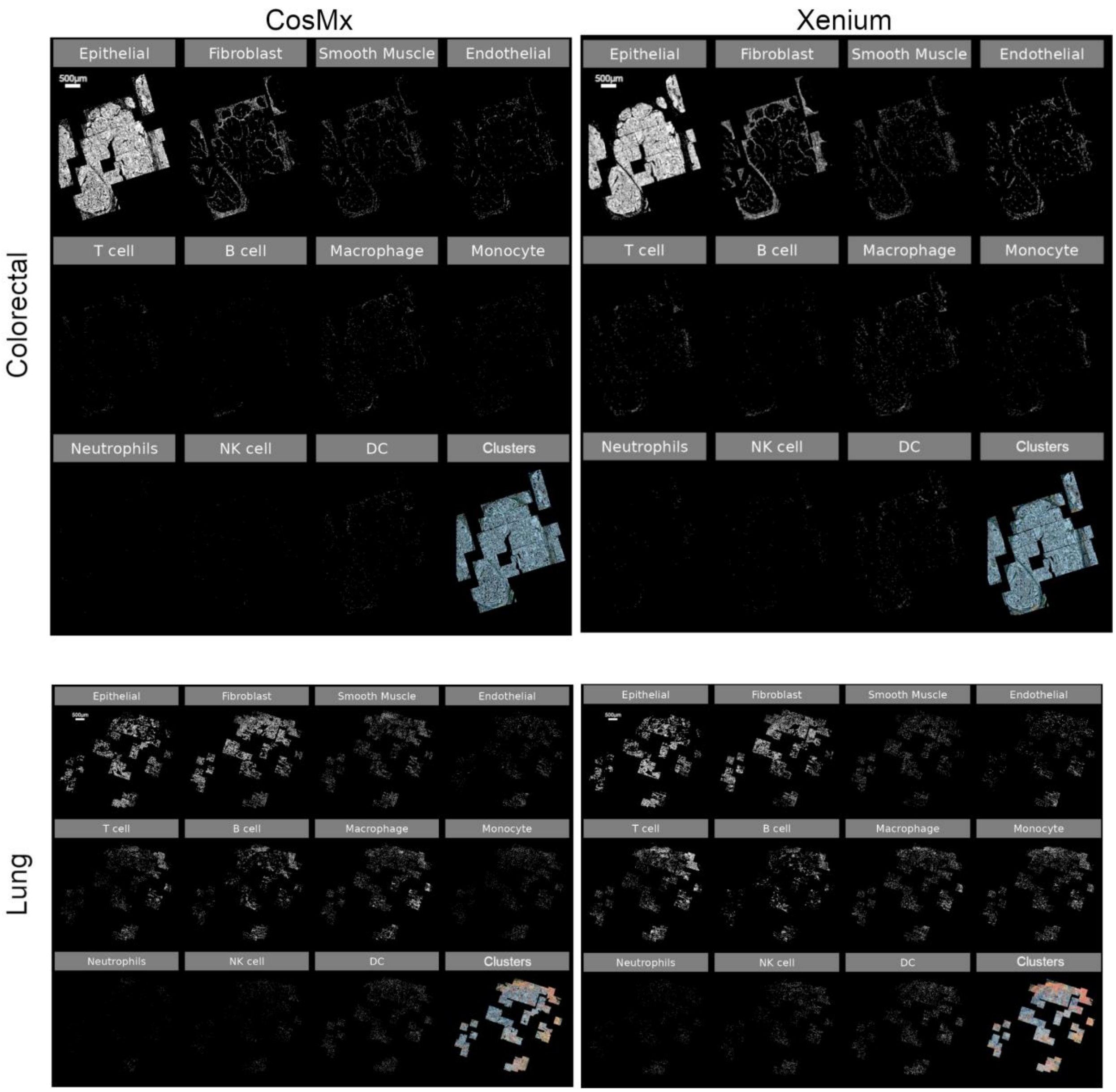

CosMx
Xenium
Colorectal
Lung
Epithelial
Fibroblast
Smooth Muscle
Endothelial
T cell
B cell
Macrophage
Monocyte
Neutrophils
NK cell
DC
Clusters
500μm

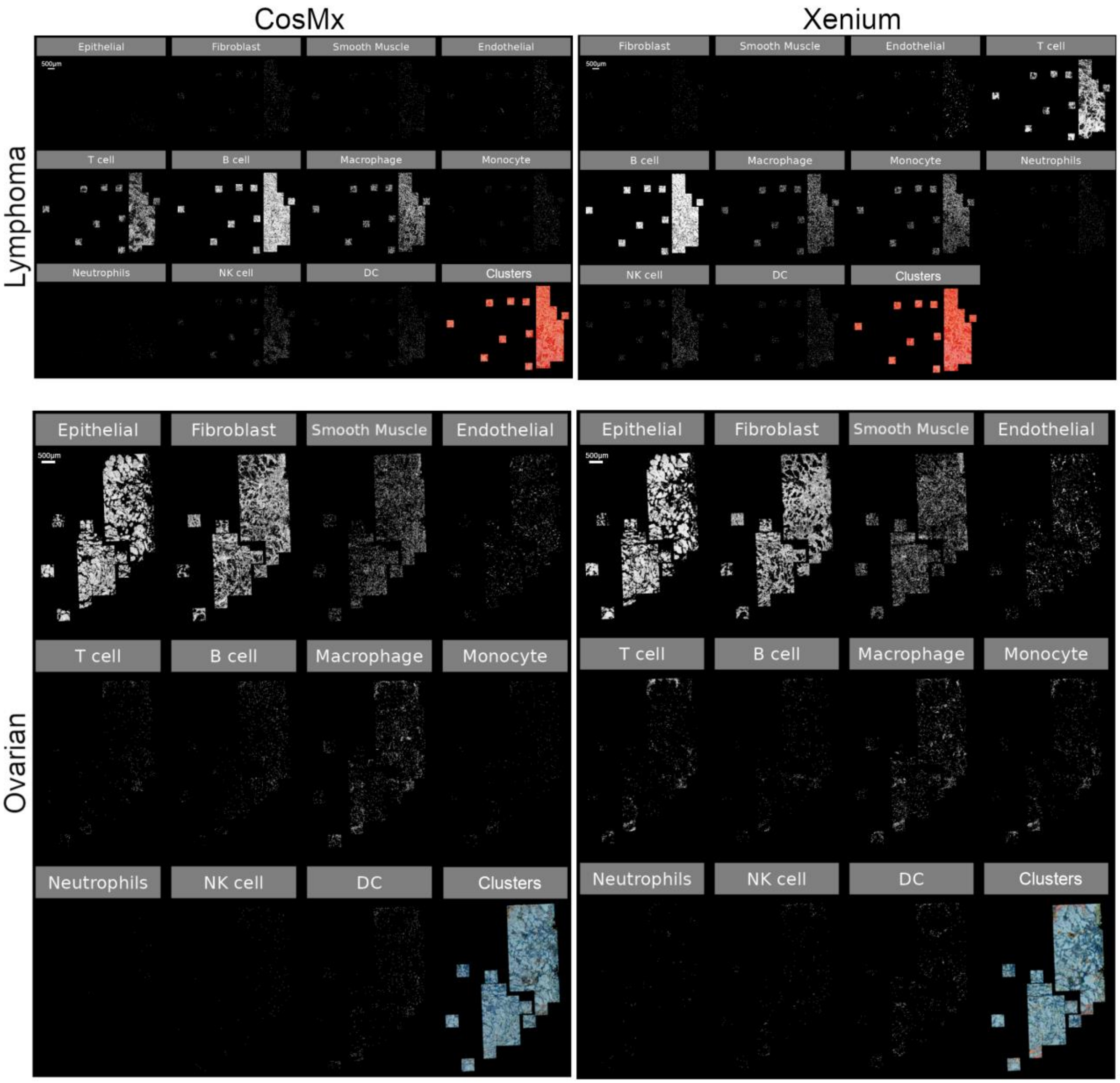


**Figure S7:** Split cell type clusters obtained from SingleR in CosMx (left) and Xenium (right) samples. The "Clusters" panel shows all cells colored by cell type.

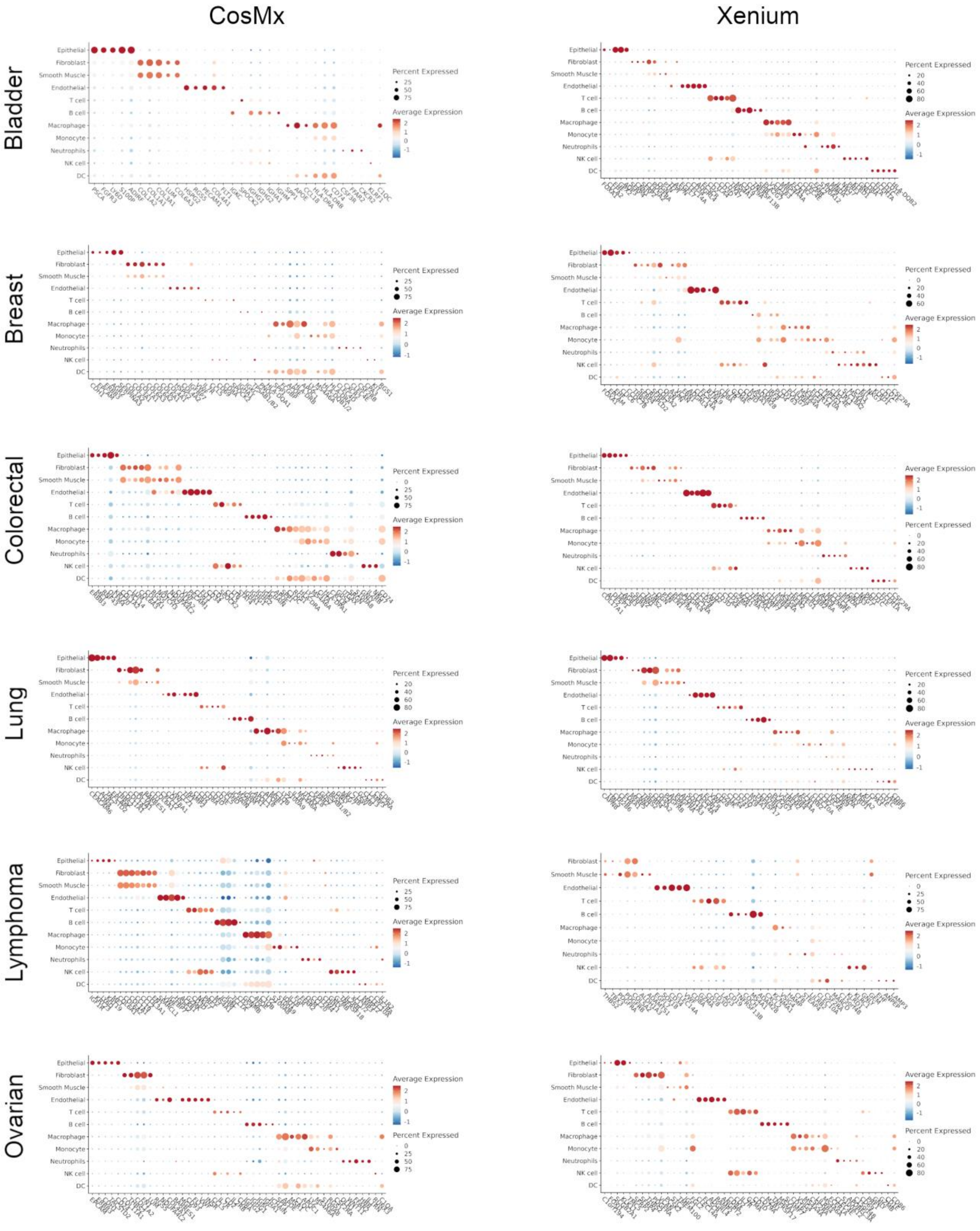


**Figure S8:** Dot plot showing scaled mean expression (dot color) and percent of expressing cells (dot size) of top 5 DEGs in each SingleR cluster.

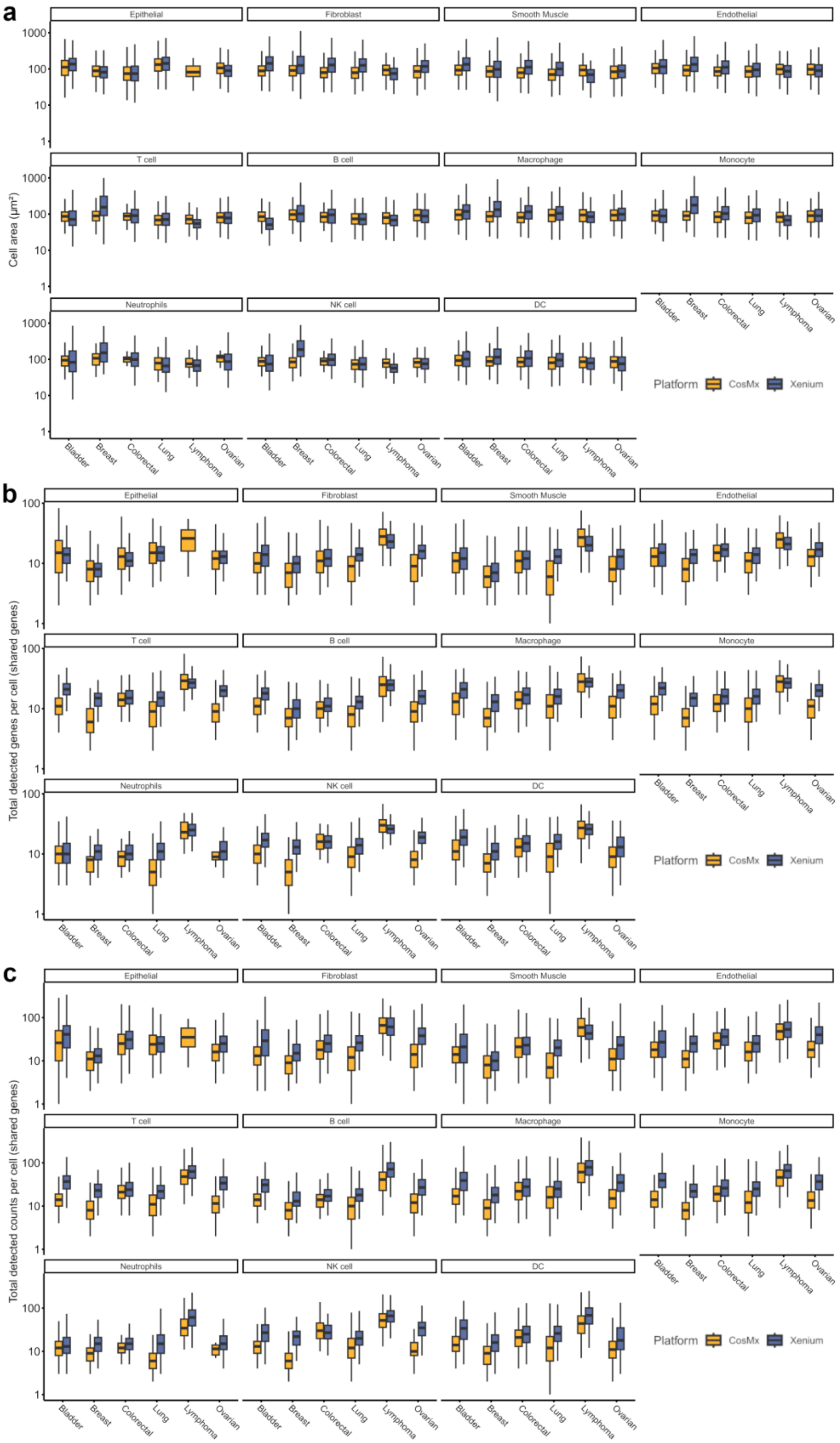
a
b
c
Epithelial
Fibroblast
Smooth Muscle
Endothelial
T cell
B cell
Macrophage
Monocyte
Neutrophils
NK cell
DC
Cell area (µm²)
Total detected genes per cell (shared genes)
Total detected counts per cell (shared genes)
1000
100
10
1
Bladder
Breast
Colorectal
Lung
Lymphoma
Ovarian
Platform
CosMx
Xenium

**Figure S9:** Boxplots showing the distribution of cell area (**a**), detected genes (**b**) and detected transcripts (**c**) in each cell in CosMx and Xenium split by cell type.

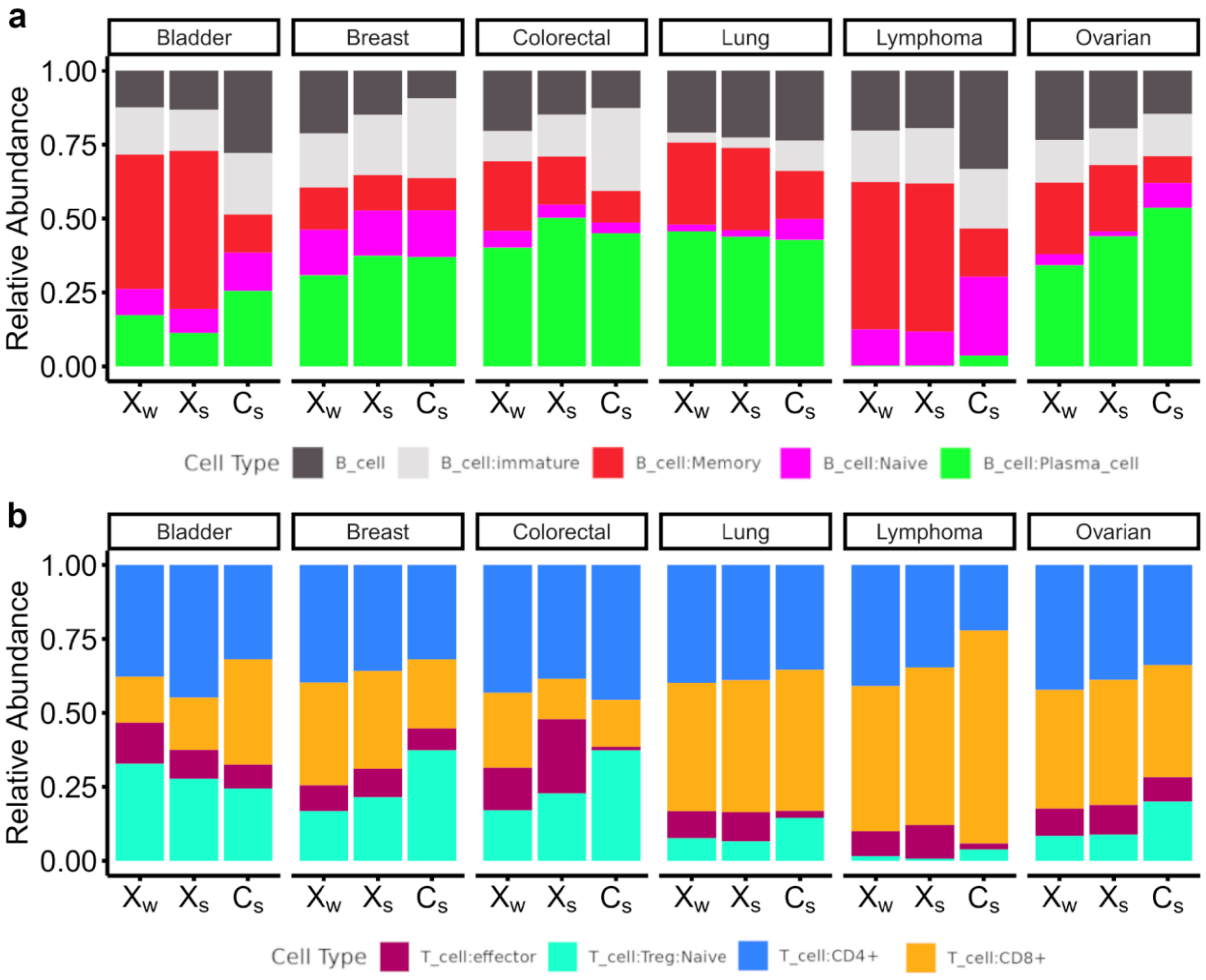


**Figure S10:** Barplot showing the relative abundance of B (**a**) and T (**b**) cell subtypes found in the whole Xenium sample(left), the common area of the Xenium cells (middle) and the common area of the CosMx cells in each tumor sample.

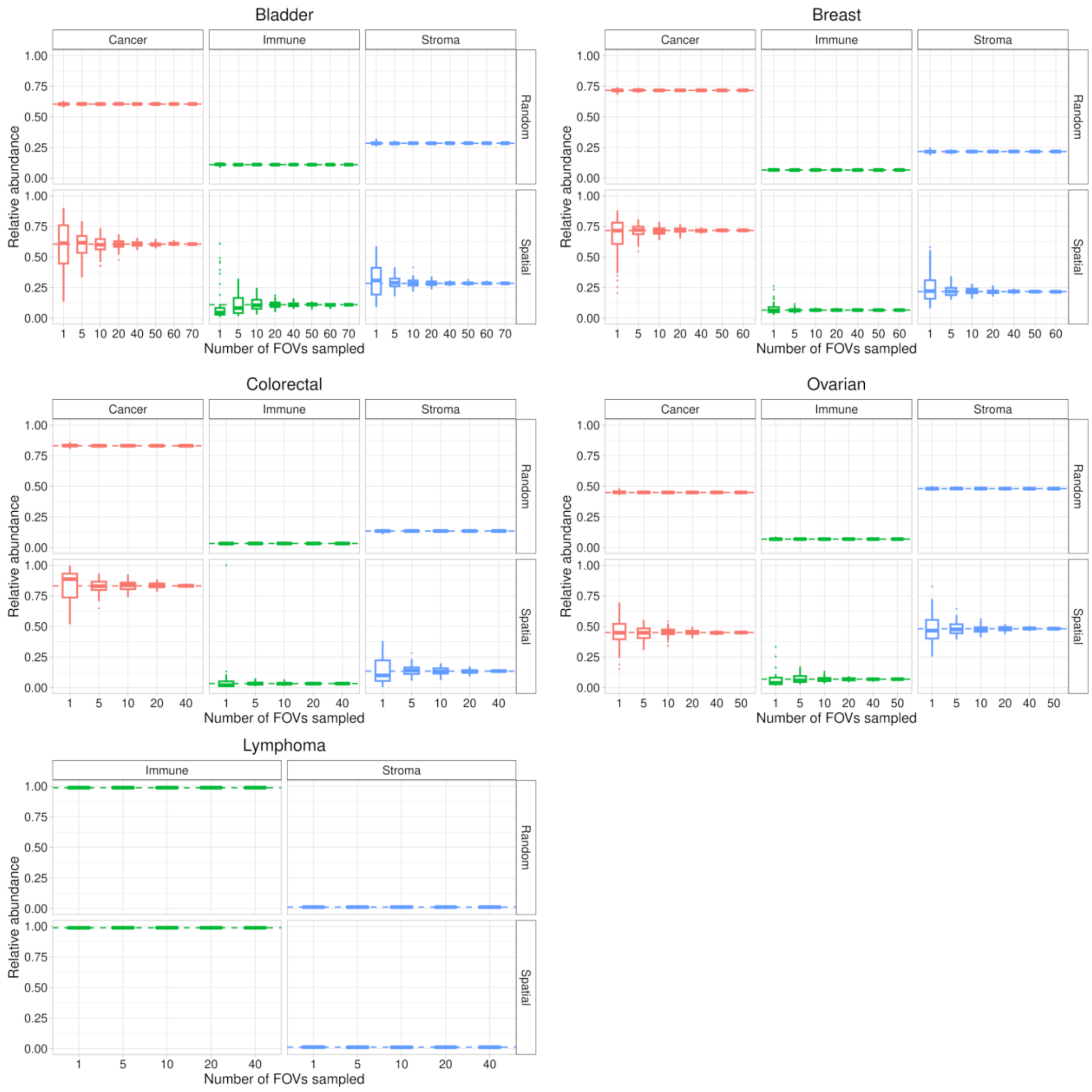


**Figure S11:** Boxplot showing the effect of sampling smaller regions to compute relative abundance in the xenium samples, of Cancer (epithelial), Stroma cells (fibroblast, smooth muscle and endothelial) and Immune (remaining cell types). In the Random category, the subset was determined by randomly sampling across the Xenium (shared area), while in the Spatial category, the sampling was based on FOV as a sampling unit. In each group number of sampled FOV, we used 100 stochastic iterations to compute the abundance of each cell group. Dashed lines represent the relative abundance of the Xenium (shared area) sample.

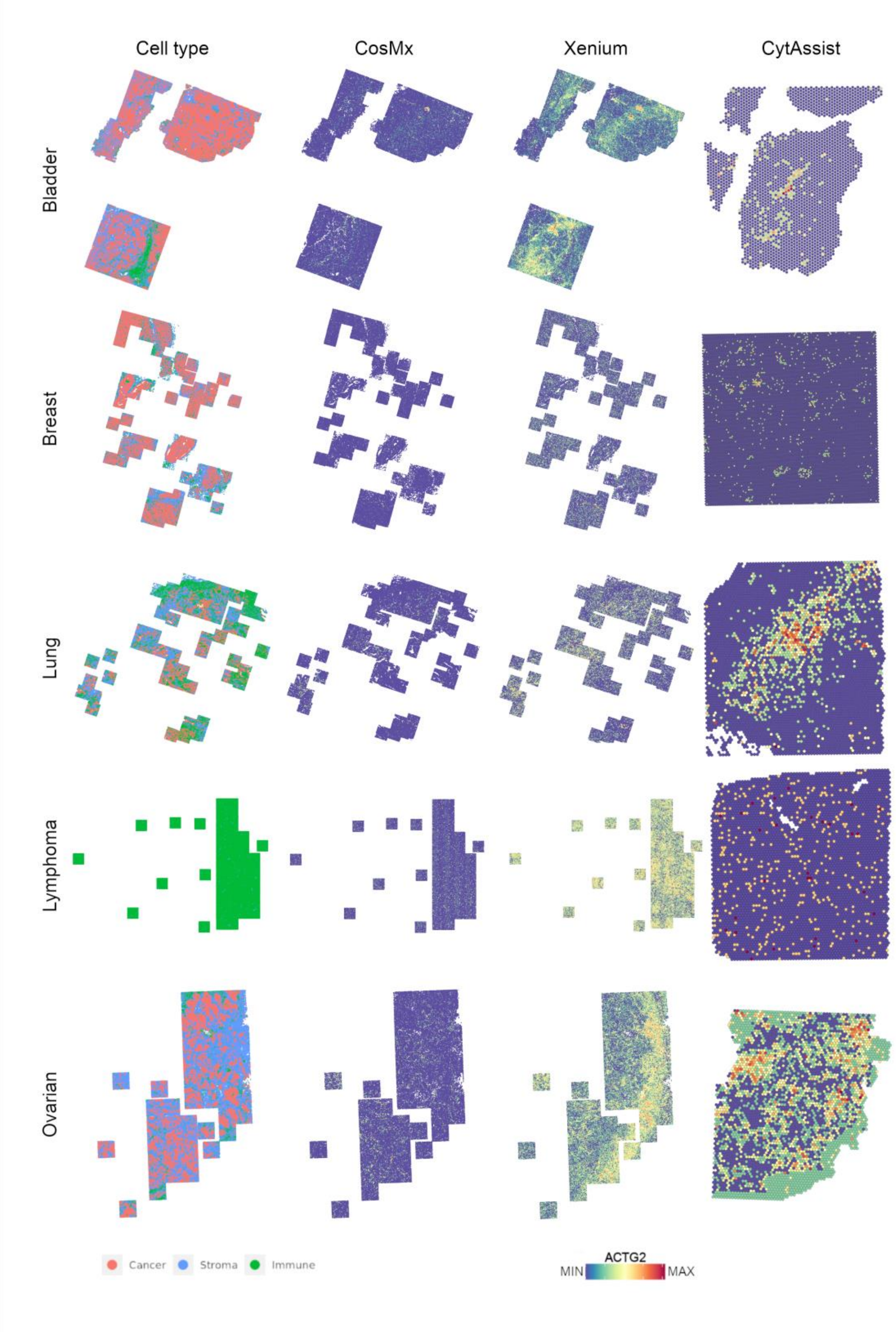


**Figure S12:** Representation of the cancer, stromal and immune cells in the Xenium sample (first) and *ACTG2* gene expression in the CosMx (second) and Xenium (third) and CytAssist (fourth) samples.

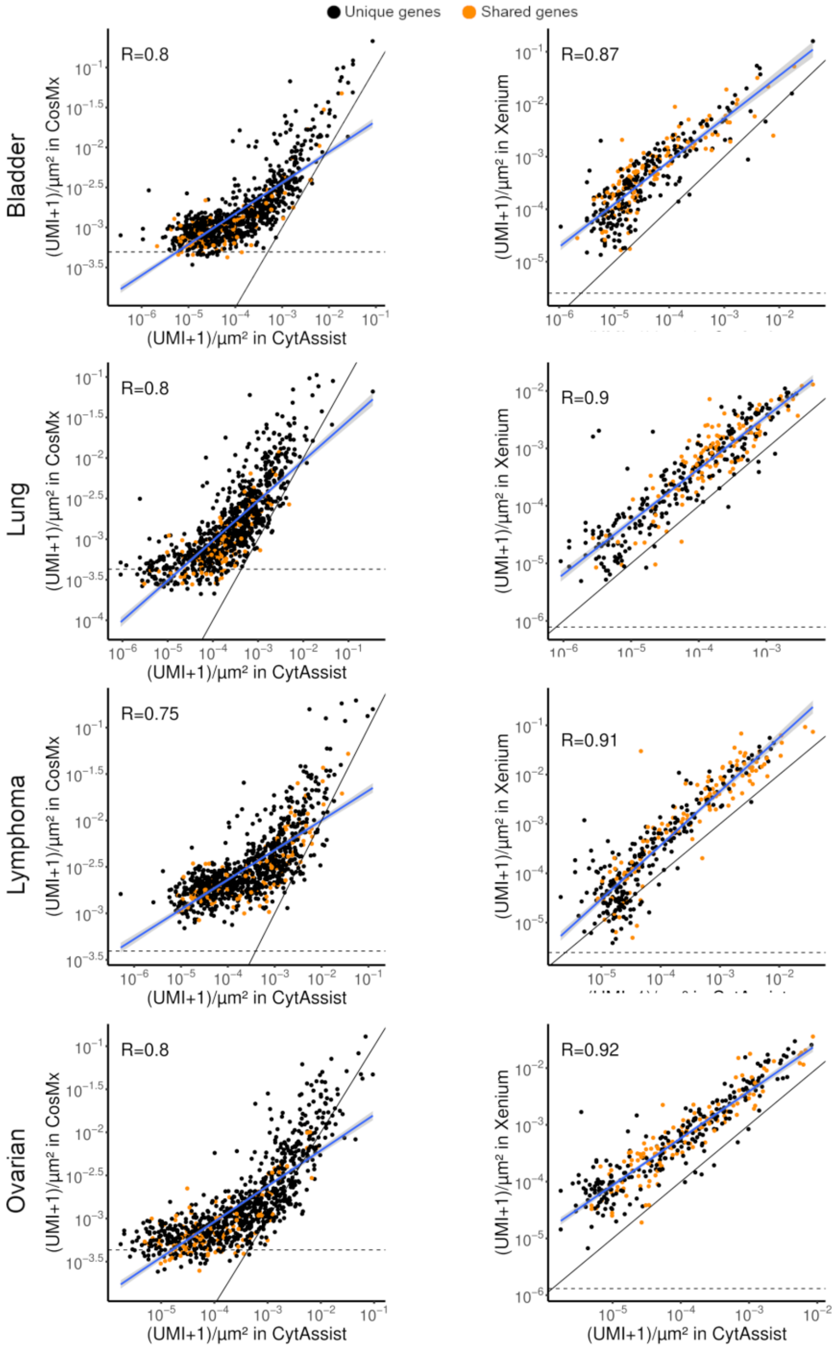


**Figure S13:** Scatterplots showing the correlation between density gene expression in CytAssist (x-axis) and CosMx (y-axis, left) or Xenium (y-axis, right). The horizontal dashed lines show the average detection rate of the negative probes in Xenium and CosMx. The diagonal dashed line shows the expected perfect correlation (i.e x = y). The blue line shows the correlation between CosMx/Xenium and CytAssist.

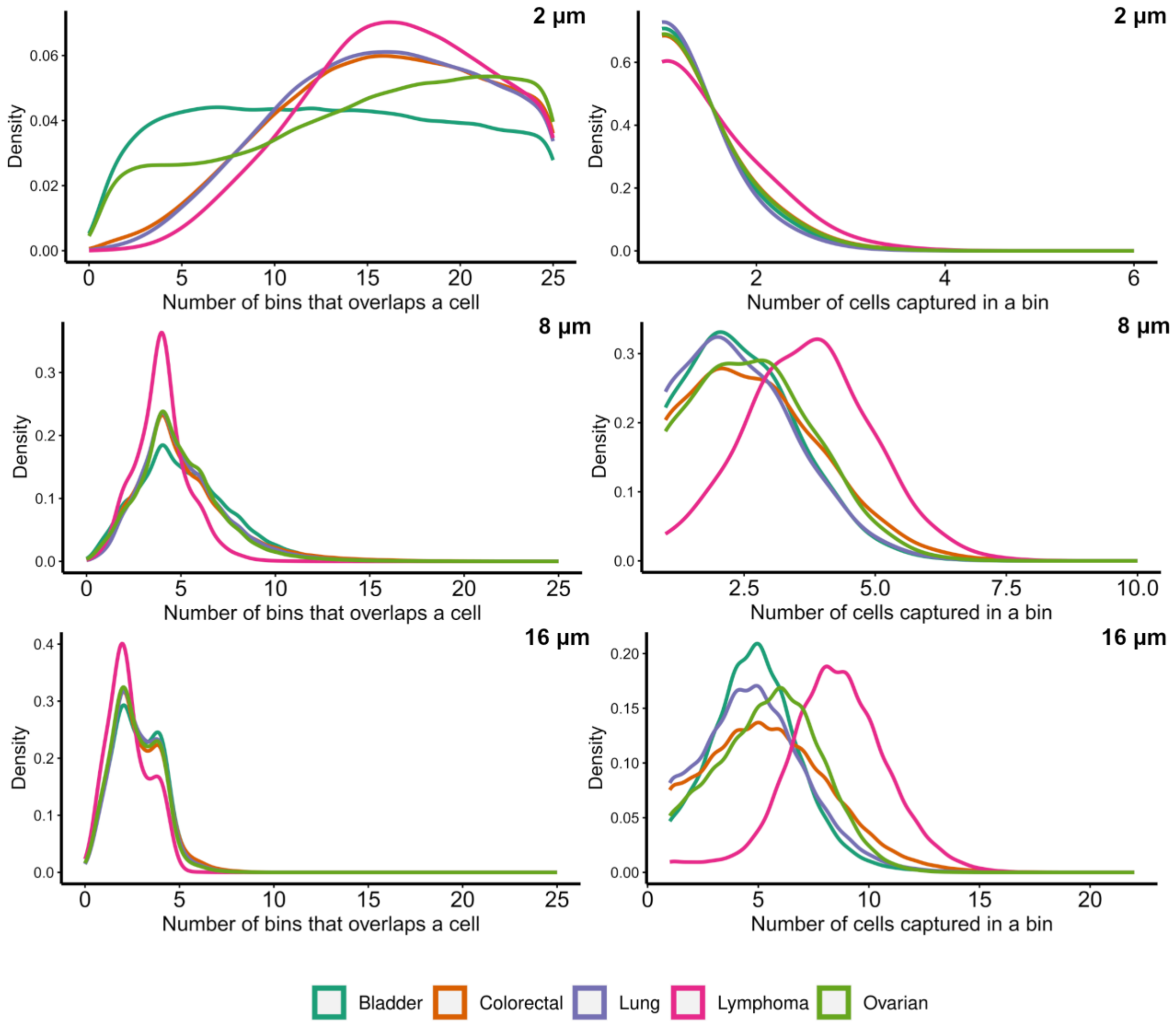


**Figure S14:** Density plot of binning simulation in Xenium samples, showing the number of different cells with transcripts in each bin split by sample (left)**.** Density plot of binning simulation in Xenium samples, showing the number of bins overlapping with transcripts from the same cell split by sample (right). Bin resolutions are indicated in the top right corner.

a

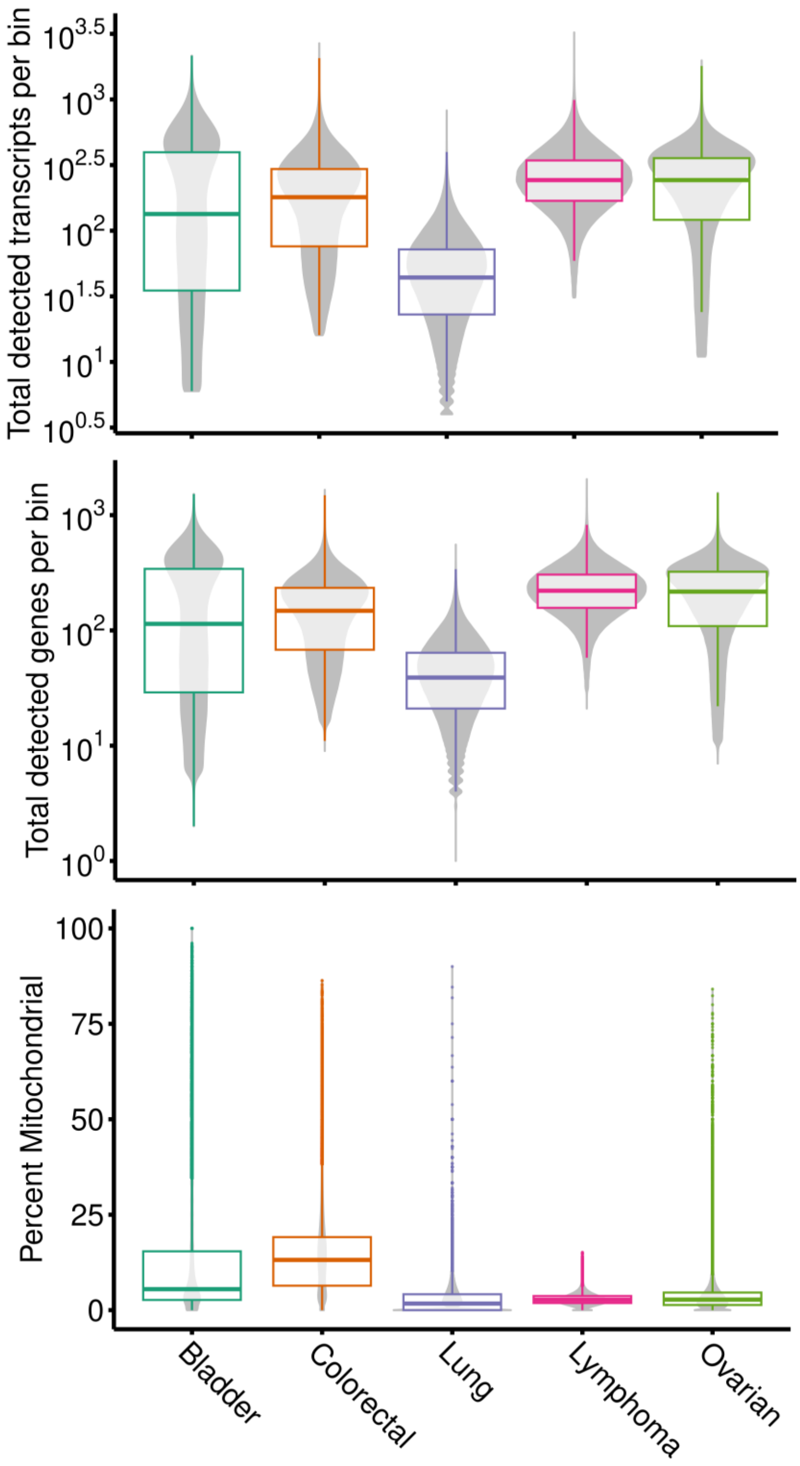

Total detected transcripts per bin
$10^{3.5}$
$10^{3}$
$10^{2.5}$
$10^{2}$
$10^{1.5}$
$10^{1}$
$10^{0.5}$
Total detected genes per bin
$10^{3}$
$10^{2}$
$10^{1}$
$10^{0}$
Percent Mitochondrial
100
75
50
25
0
Bladder
Colorectal
Lung
Lymphoma
Ovarian

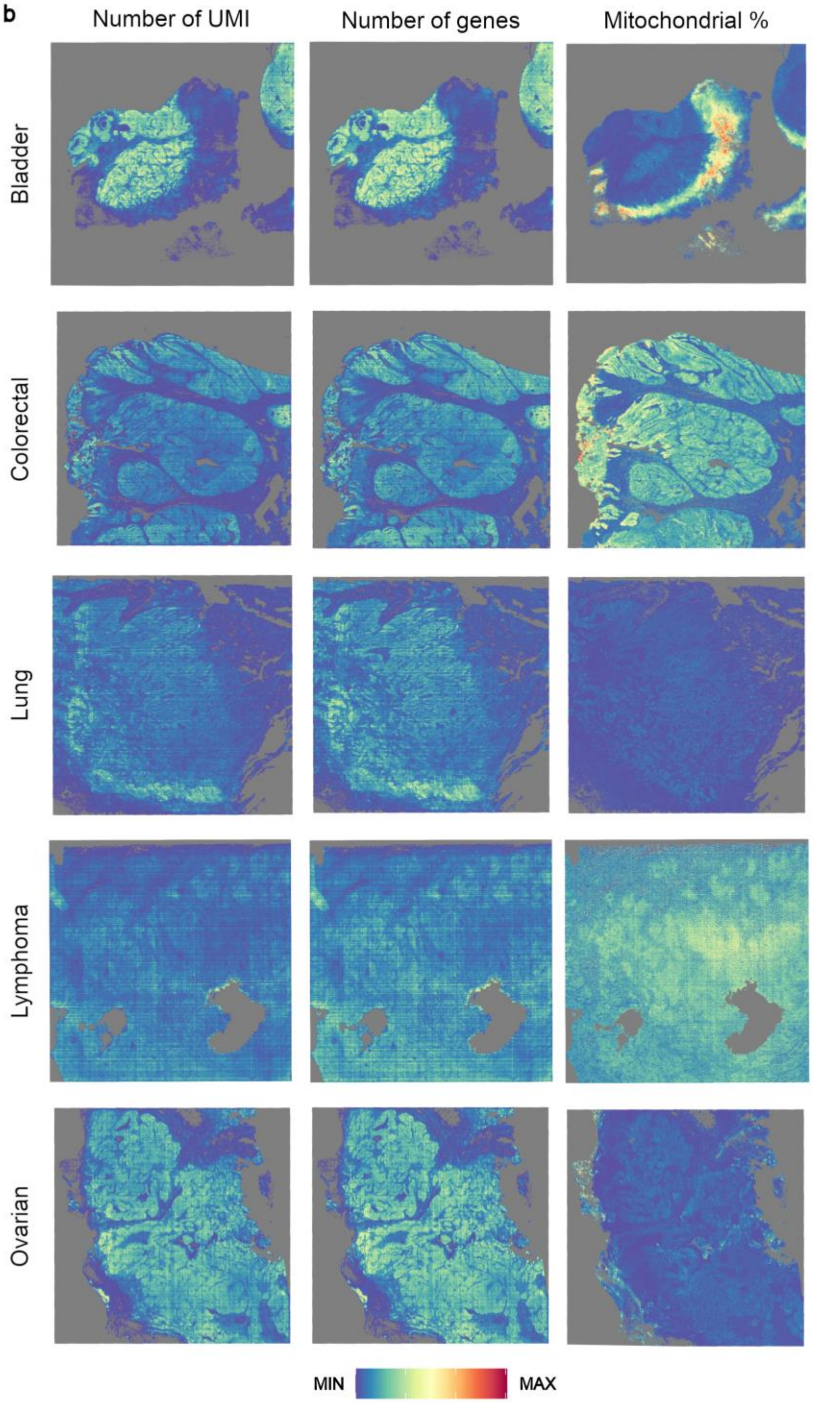


**Figure S15: a)** Distribution of the number of transcripts (top), genes (middle) and mitochondrial percentage (bottom) detected in each 8 µm bin across samples**. b)** Spatial expression of the number of transcripts (left), genes (middle) and mitochondrial percentage (right) in each 8 µm bin. Ranges of each QC are the same as a).

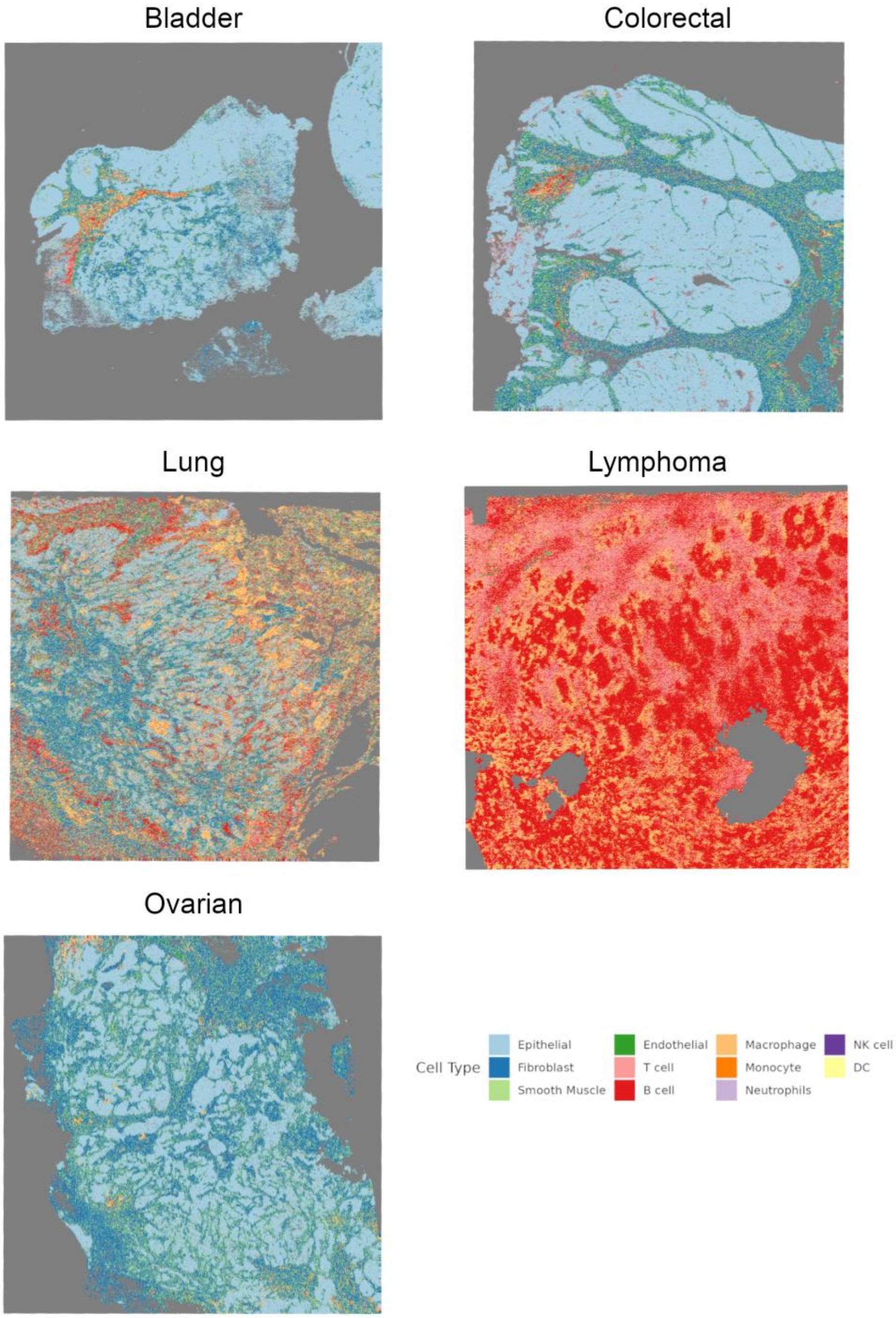


**Figure S16:** Cell type clusters obtained from SingleR in VisiumHD samples.

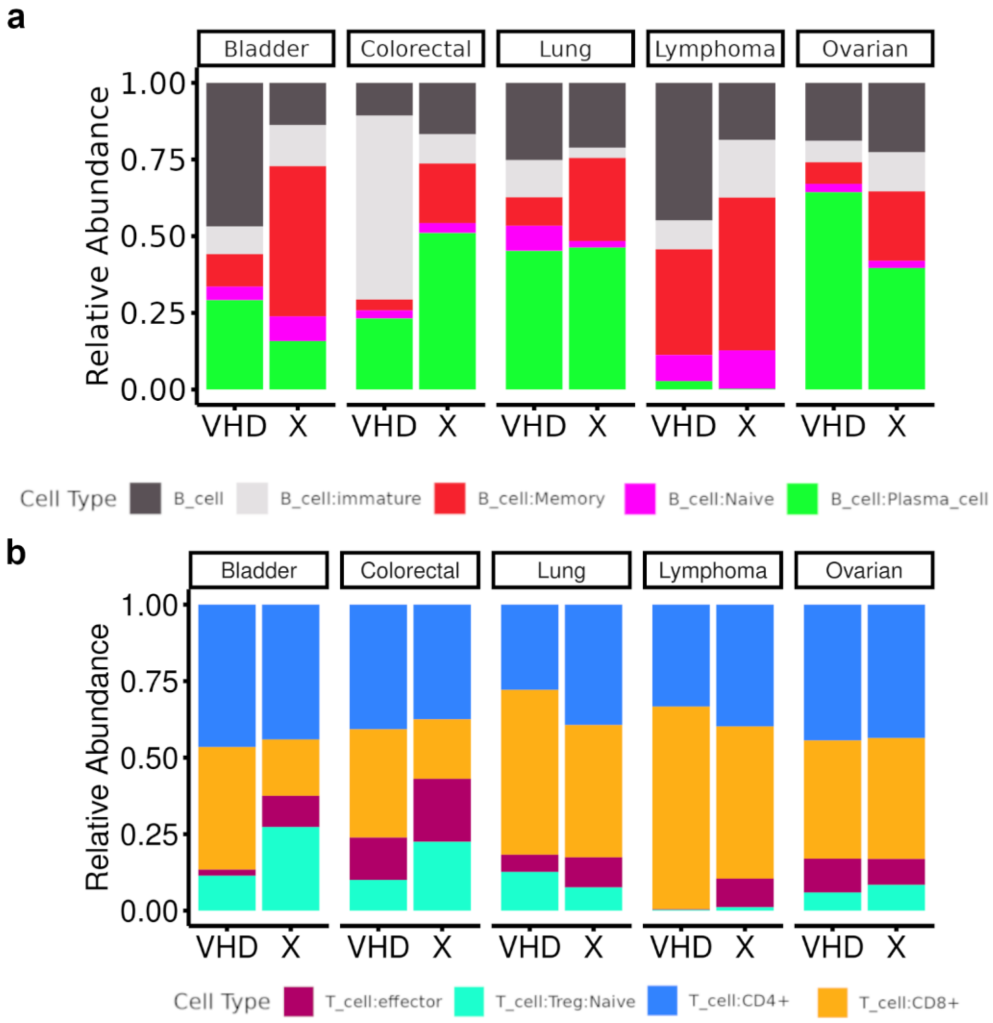


**Figure S17:** Barplot showing the relative abundance of B (**a**) and T (**b**) cell subtypes found in the VisiumHD (VHD) and Xenium (X) in each tumor sample.

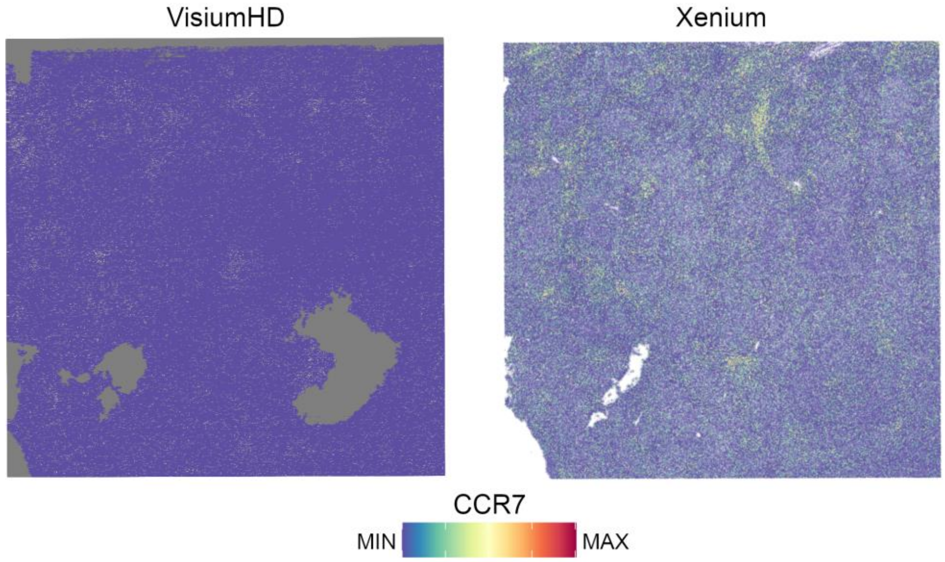


**Figure S18:** Plot showing the expression of *CCR7* in the lymphoma sample with VisiumHD (left) and Xenium (right).
